# COP9 signalosome component CSN-5 stabilizes PUF proteins FBF-1 and FBF-2 in *Caenorhabditis elegans* germline stem cells

**DOI:** 10.1101/2022.06.22.497213

**Authors:** Emily Osterli, Mary Ellenbecker, Xiaobo Wang, Mikaya Terzo, Ketch Jacobson, DeAnna Cuello, Ekaterina Voronina

**Author notes:** current address: Dermatology Hospital, Southern Medical University, Guangzhou, China (Xiaobo Wang).

## Abstract

RNA-binding proteins FBF-1 and FBF-2 (FBFs) are required for germline stem cell maintenance and the sperm/oocyte switch in *Caenorhabditis elegans*, though the mechanisms controlling FBF protein levels remain unknown. We identified an interaction between both FBFs and CSN-5, a component of the COP9 (constitutive photomorphogenesis 9) signalosome. Here, we find that the MPN (Mpr1/Pad1 N terminal) metalloprotease domain of CSN-5 interacts with the PUF (Pumilio and FBF) RNA-binding domain of FBFs and the interaction is conserved for human homologs PUM1 and CSN5. The interaction between FBF-2 and CSN-5 can be detected *in vivo* by proximity ligation. *csn-5* mutation results in destabilization of FBF proteins, a decrease in the numbers of germline stem cells, and disruption of the switch from spermatogenesis to oogenesis. The loss of *csn-5* does not decrease the levels of a related PUF protein PUF-3 and *csn-5(lf*) phenotype is not enhanced by *fbf-1/2* depletion, suggesting that the effect is specific to FBFs. The effect of *csn-5* on germline sex determination is largely independent of the COP9 signalosome and is cell autonomous. Surprisingly, regulation of FBF protein levels involves a combination of COP9-dependent and –independent mechanisms differentially affecting FBF-1 and FBF-2. This work supports a previously unappreciated role for CSN-5 in stabilization of germline stem cell regulatory proteins FBF-1 and FBF-2.

**Author Summary:** Germ cell development and reproductive success in the nematode *C. elegans* rely on the function of germline stem cells. Continued maintenance of these cells is supported by the activity of conserved RNA-binding proteins FBF-1 and FBF-2 (FBFs). However, it is unknown how FBF protein levels are regulated. Here, we identify a direct interaction between FBFs and CSN-5, a component of the COP9 signalosome best known for its role in regulating protein degradation. We find that CSN-5 promotes FBF stability and allows for accumulation of steady-state protein levels, thereby promoting FBF function. In *csn-5* mutants, we find a significant reduction of FBF proteins, decrease of stem cells, and failure to promote oogenesis consistent with compromised FBF function. Furthermore, CSN-5 contributes to FBF protein stability through two mechanisms. This work demonstrates a previously unappreciated role for CSN-5 in stabilization of FBF proteins. Based on our finding that the FBF/CSN-5 interaction is conserved and detectable between homologous human proteins, we speculate this relationship might be relevant for understanding stem cell maintenance in a range of species, from nematodes to humans.

## Introduction

Stem cells are a population of unspecialized cells with the capability to both self-renew and preserve the stem cell pool or to differentiate and support tissue maintenance (Morrison and Kimble, 2006). Stem cell function depends on faithful expression of essential developmental regulators (Morrison et al., 1997). A number of mechanisms controlling expression of stem cell machinery at the levels of epigenetics, transcription, and RNA stability have been widely studied (for example, Lee et al., 2016; Chen et al., 2020; Samuels et al., 2020; Demarco et al., 2022; Vogiatzoglou et al., 2022). As the stem cell proteome is exquisitely controlled in both self-renewing state and through developmental transitions (Vilchez et al., 2014; Llamas et al., 2020), it is critical to understand the mechanisms regulating stability of key self-renewal factors.

We use the *C. elegans* germline to study stem cell regulators FBF-1 and FBF-2 (FBFs). The *C. elegans* gonad consists of two U-shaped arms, each with a population of stem and progenitor cells (SPCs) at the distal end (Fig. 1A; Kimble and White, 1981). SPCs are maintained through the GLP-1/Notch signaling from the somatic niche cell (Austin and Kimble, 1987; Berry et al., 1997; Kimble and Crittenden, 2007). As the cells progress away from the niche, they begin to differentiate and enter meiosis before generating gametes at the proximal end (Hubbard, 2007; Pazdernik and Schedl, 2013; Voronina and Greenstein, 2016). The hermaphrodite germlines produce sperm during larval development and switch to oogenesis in the young adults (Pazdernik and Schedl, 2013).

**Fig. 1.**
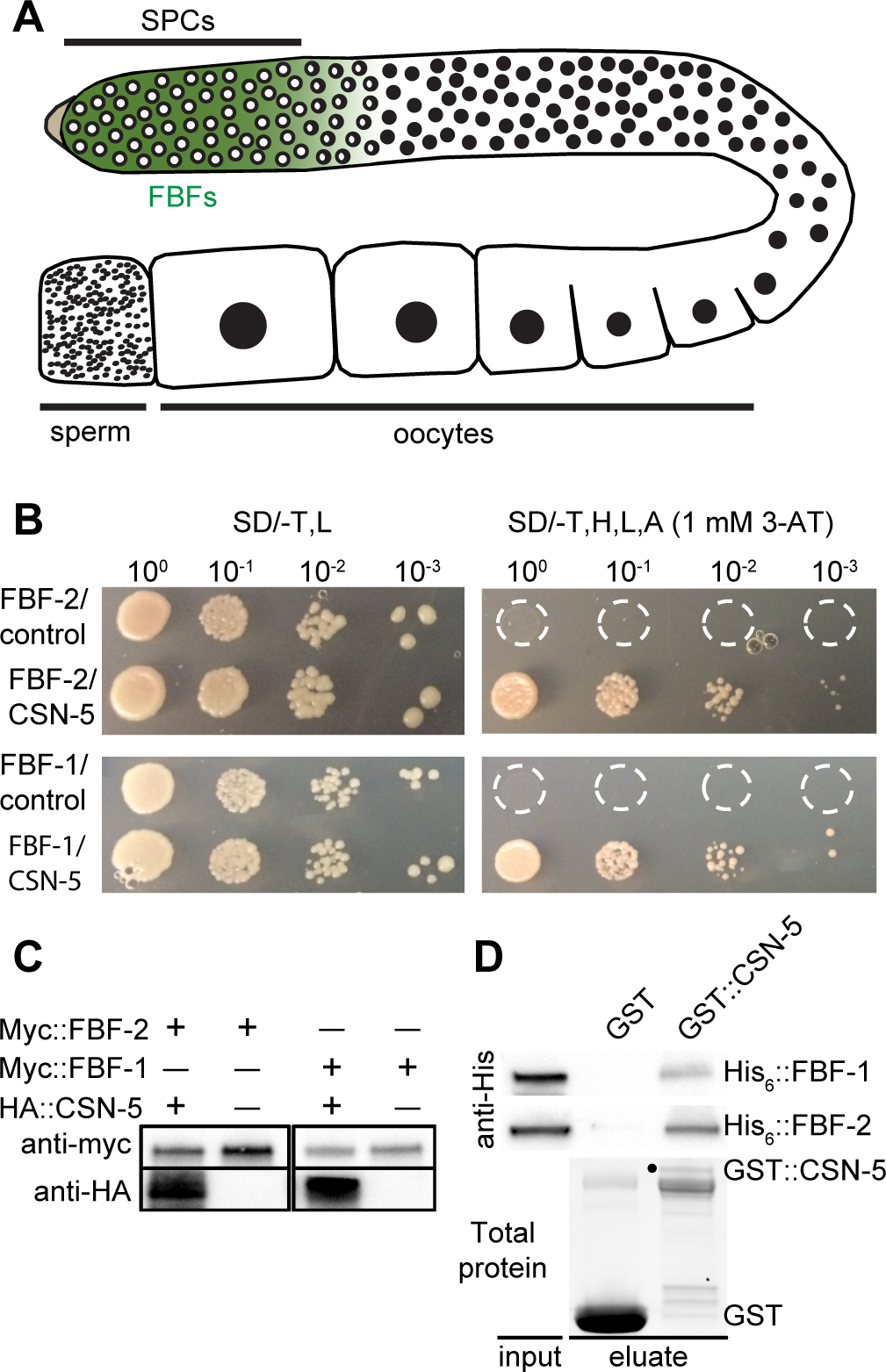
CSN-5 interacts with FBFs. (A) Schematic of adult *C. elegans* hermaphrodite gonad. Stem and progenitor cells (SPCs) expressing FBF proteins are highlighted in green. COP9 subunit CSN-5 is expressed throughout oogenic germline. (B) Interaction between CSN-5 and FBF-1/-2 is detected in a yeast two-hybrid assay with FBFs fused to Gal4 DNA-binding domain as baits and CSN-5 fused to Gal4 activation domain as prey. This experiment was performed in two replicates. (C) Western blot confirming expression of FBFs and CSN-5 in yeast cultures. (D) GST pulldown assay detects a direct interaction between GST::CSN-5 and both His_6_::FBF-1 and His_6_::FBF-2. FBFs are detected by Western blot with anti-His. Total protein (representative image from FBF-2 pulldown) detected by stain-free chemistry, the position of GST::CSN-5 is marked with a black dot.

FBFs are RNA-binding proteins belonging to the highly conserved PUF (Pumilio and FBF) family (Wickens et al., 2002). FBFs are enriched in SPCs (Fig. 1A) and act by binding to and repressing their target mRNAs (Zhang et al., 1997; Crittenden et al., 2002; Prasad et al., 2016). FBFs promote germline stem cell maintenance, and *fbf-1 fbf-2* double mutant germline stem cells prematurely enter meiosis during late larval development (Crittenden et al., 2002; Wickens et al., 2002; Wang and Voronina, 2020). Consistent with this function, many FBF targets encode developmental regulators, which are silenced in stem cells and then activated upon differentiation, such as the differentiation-promoting protein GLD-1 (Crittenden et al., 2002; Suh et al., 2009). FBFs also promote self-renewal of germline stem cells by repressing *cki-2* (Kalchhauser et al., 2011), a Cyclin E/Cdk2 inhibitor which regulates the decision to enter/exit the cell cycle (Buck et al., 2009). Additionally, FBF-1 and FBF-2 regulate germline sex determination by facilitating the switch from spermatogenesis to oogenesis (Zhang et al., 1997) as *fbf-1 fbf-2* double mutant gonads fail to undergo oogenesis, resulting in a masculinization of germline (Mog) phenotype where only sperm is produced (Crittenden et al., 2002). Despite these well-characterized functions of FBFs, the post-translational mechanisms regulating the levels of FBF proteins in stem cells remain unknown.

One prominent cellular mechanism impacting protein stability and steady-state levels involves the COP9 (constitutive photomorphogenesis 9) signalosome, a highly conserved enzymatic complex composed of eight subunits, denoted CSN1 to CSN8 (Qin et al., 2020). Although initially discovered for its role in regulation of transcription (Wei and Deng, 1992), the COP9 signalosome’s best documented function is as a regulator of protein degradation (Wei et al., 1994; Chamovitz et al., 1996; Claret et al., 1996; Chamovitz, 2009). COP9 impacts protein degradation by removing a ubiquitin-like protein, NEDD8, from cullin subunits of E3 ubiquitin ligases (Cope et al., 2002; Lingaraju et al., 2014). The balance between neddylation and deneddylation is essential to maintain the activity of cullin-based ubiquitin ligases and to facilitate the degradation of their substrates (Lyapina et al., 2001; Doronkin et al., 2003; Pintard et al., 2003; Wu et al., 2005). The overall architecture of COP9 resembles the 19S lid of the 26S proteasome, which is responsible for the majority of intracellular protein degradation (Enchev et al., 2012; Lingaraju et al., 2014). CSN subunits 1-4, 7, and 8 contain PCI (**P**roteasome, **C**OP9 signalosome, translation **I**nitiation factor) domains similar to the six subunits of the proteasome lid (Dessau et al., 2008; Hofmann and Bucher, 1998). COP9 signalosome subunit 5 (CSN5) is the only catalytically competent component and contains a JAB1/MPN (**M**pr1/**P**ad1 **N**-terminal) metalloprotease domain, which is responsible for the complex’s deneddylating activity (Zhang et al., 2012; Echalier et al., 2013). The signalosome is inactive without the incorporation of CSN5, and likewise, the current literature suggests CSN5 is inactive on its own and unable to deneddylate cullins in isolation (Cope et al., 2002; Sharon et al., 2009; Echalier et al., 2013; Lingaraju et al., 2014). Furthermore, CSN5 must dimerize with the other MPN domain containing COP9 subunit CSN6, to be incorporated into the COP9 holoenzyme and become activated (Birol et al., 2014; Lingaraju et al., 2014). Apart from COP9 subunits, CSN5 has been reported to interact with other cellular proteins (Tomoda et al., 1999; Smith et al., 2002; Yoshida et al., 2013; Shackleford and Claret, 2010) and to promote accumulation of several of its partners (Bae et al., 2002; Bemis et al., 2004; Orsborn et al., 2007; Liu et al., 2009; Lim et al., 2016). Furthermore, CSN5 interaction partners may be stabilized in a manner dependent on the entire COP9 complex (Wu et al., 2009) or independently of the COP9 holoenzyme (Bemis et al., 2004).

In a yeast-two hybrid screen for FBF binding partners, we identified *C. elegans* CSN-5 as an interactor of FBF-2. This work investigates CSN-5’s contribution to FBF function in the germline. We find that CSN-5 stabilizes both FBF-1 and FBF-2 in germline SPCs, correlating with effective stem cell maintenance and female fate. Our findings identify a previously unappreciated role for CSN-5 and we speculate its contribution to stem cell regulatory protein accumulation is relevant to a wide range of organisms.

## Results

### CSN-5 is a new interacting partner of FBFs

To identify proteins that associate with FBF-2, we performed a yeast two-hybrid screen of a *C. elegans* cDNA library with FBF-2 as bait (see Materials and Methods for detail). Briefly, the yeast transformed with full-length FBF-2 fused to the Gal4 DNA-binding domain were mated to a yeast library of mixed-stage wild type *C. elegans* cDNAs fused to the Gal4 activation domain. Resulting diploids were selected for growth on minimal media lacking histidine and adenine selecting for expression of *HIS3* and *ADE2*, with the addition of 1 mM 3-AT (a competitive inhibitor of the HIS3 enzyme) added for stringency. The screen of estimated 2x10^8^ yeast diploids identified a known FBF interactor SYGL-1 (Shin et al., 2017) as well as several potential new interactors (Table S1), with one of the most abundant partners representing a fragment of COP9 signalosome subunit CSN-5. To validate this potential interaction and to test whether CSN-5 might bind FBF-1 as well as FBF-2, we performed a directed yeast two-hybrid assay, in which full-length FBF-1 or FBF-2 fused to the Gal4 DNA-binding domain were cotransformed with full-length CSN-5 fused to the Gal4 activation domain. Robust growth was observed when CSN-5 was co-transformed with either FBF-1 or FBF-2, but not in controls (Fig. 1B). Yeast expression of c-myc-tagged FBFs and HA-tagged CSN-5 was detected by Western blot using anti-Myc and anti-HA, respectively (Fig. 1C). We conclude that CSN-5 interacts with both FBF-1 and FBF-2 in yeast. Additionally, we performed a GST pulldown assay with bacterially-expressed GST-tagged CSN-5 and His_6_-tagged FBFs, which detected direct interaction between these proteins (Fig. 1D).

### The interaction between the RNA-binding domain of FBFs and the MPN domain of CSN-5 is conserved in evolution

To elucidate the location of the interacting domains within FBFs and CSN-5, we generated several truncation constructs of FBF-1, FBF-2, and CSN-5 expressing distinct domains amenable to recombinant expression based on the previous studies (Fig. 2B, D; Bernstein et al., 2005; Echalier et al., 2013; Birol et al., 2014), and tested their association via GST pulldown assay. We observed that the conserved RNA binding domains (RBDs) of both FBFs were sufficient for the interaction with CSN-5, and the binding between FBF-2 and CSN-5 was lost in the absence of FBF-2 RBD (Fig. 2A). Furthermore, the interaction between FBF-2 and CSN-5 was dependent on CSN-5^MPN^ domain (Fig. 2C). We then tested whether FBF-2 could interact with the other MPN domain of COP9 signalosome, contributed by CSN-6. The MPN domain of CSN-6 was delineated by homology to human CSN6 (Ma et al., 2014). GST pulldown assay suggested that the interaction of His_6_::FBF-2 with GST::CSN-6^MPN^ is much weaker than that with CSN-5^MPN^ (Fig. 2E). We concluded that the FBF/CSN-5 interaction is mediated by the conserved, structured domains, with selectivity towards specific members of the domain families. We further tested whether FBF-2 interaction with MPN domains of CSN-5 and CSN-6 was RNA-dependent, and found that the binding was not disrupted by the RNase A treatment (Fig. S1).

**Fig. 2.**
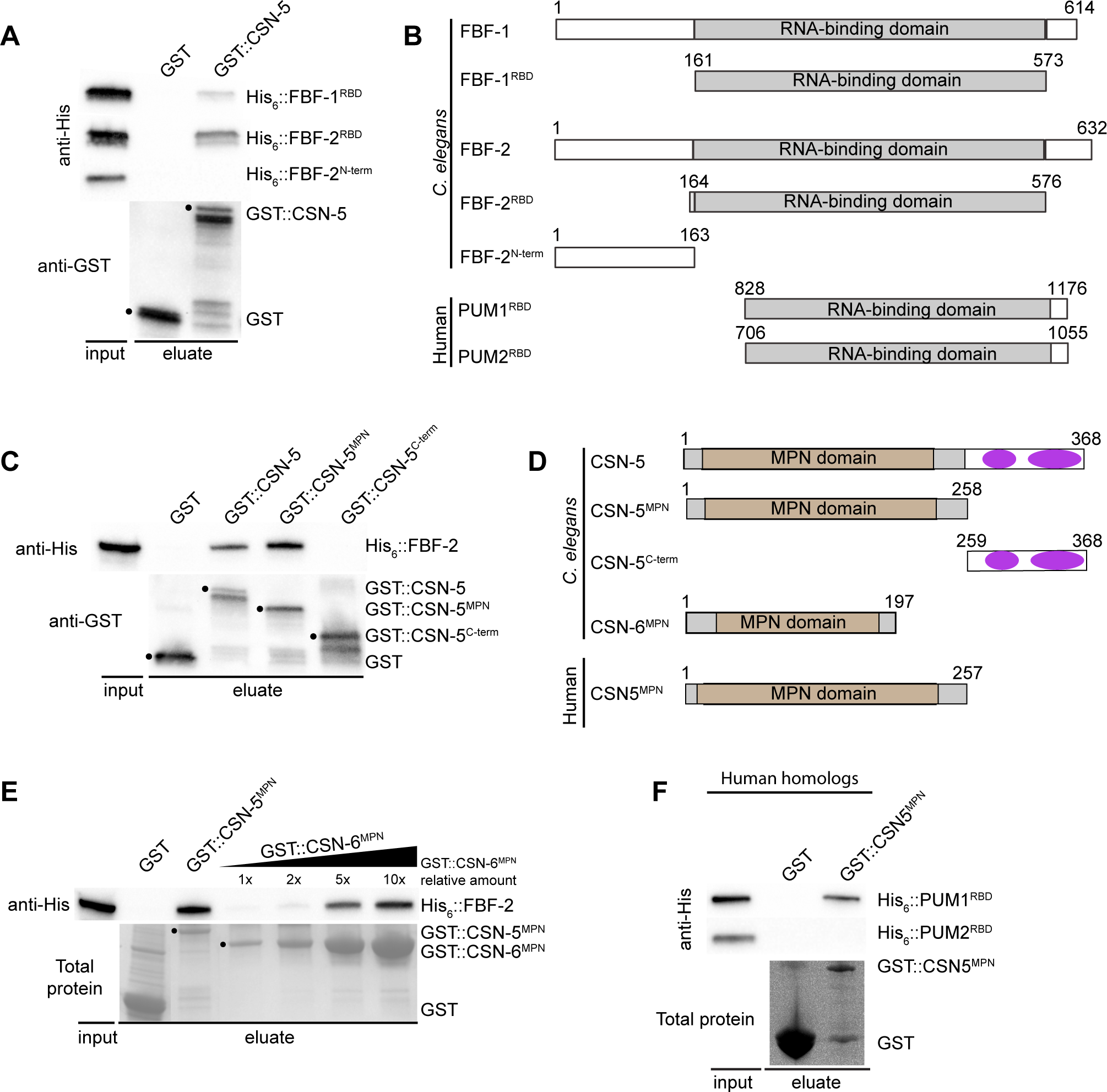
Interaction between FBF RNA-binding domain and MPN domain of CSN-5 is conserved in evolution. (A) GST or GST::CSN-5 were tested for binding His_6_::FBF-1^RBD^, His_6_::FBF-2^RBD^, and His_6_::FBF-2^N-term^ constructs. Proteins are detected by Western blot with anti-His and anti-GST, the position of GST::CSN-5 (representative image from His_6_::FBF-2^N-term^ pulldown) is marked with a black dot. (B) FBF-1/-2 and PUM1/2 truncation constructs. Grey: RNA-binding domain. Amino acid positions of the truncations are indicated above each construct. (C) GST::CSN-5 truncation constructs were tested for binding His_6_::FBF-2. Proteins are detected by Western blot with anti-His and anti-GST, the positions of GST-tagged constructs are marked with a black dot. (D) CSN-5, CSN-6, and CSN5 truncation constructs. Brown: core MPN domain. C-terminal helices (purple ovals) that incorporate into the helical bundle of the COP9 signalosome (Lingaraju et al., 2014). Amino acid positions of the truncations are indicated above each construct. (E) GST pulldown of His_6_::FBF-2 with GST::CSN-5^MPN^ and increasing concentrations of GST::CSN-6^MPN^ as detected by Western blot with anti-His. Total protein seen by Coomassie. See also Fig. S1. (F) GST pulldown of human homologs, GST::CSN5^MPN^ with His_6_::PUM1^RBD^ or His_6_::PUM2^RBD^, as detected by Western blot with anti-His. Total protein (representative image from His_6_::PUM2^RBD^ pulldown) detected by stain-free chemistry.

Both FBFs and CSN-5 have homologous human proteins, PUM1/PUM2 and CSN5, respectively (Spassov and Jurecic, 2002; Qin et al., 2020). We tested whether the RBDs and MPN domain of the corresponding human homologs (Fig. 2B, D) were able to interact. A GST pulldown assay of GST::CSN5^MPN^ with His_6_::PUM1^RBD^ and His_6_::PUM2^RBD^ revealed that the interaction we observed for the nematode proteins is conserved for human homologs CSN5 and PUM1, but not PUM2, suggesting an evolutionarily conserved protein complex (Fig. 2F). We considered whether CSN-5 were able to interact with other *C. elegans* PUF proteins including PUF-8, PUF-11, and PUF-3, but were unable to produce soluble recombinant PUF proteins to test this hypothesis.

### FBF-2 interacts with CSN-5 *in vivo*

After characterizing the interaction between FBFs and CSN-5 *in vitro*, we tested if this interaction is also observed *in vivo*. We utilized proximity ligation assays (PLA) to detect *in situ* protein-protein interactions at distances <40 nm (Söderberg et al., 2008; Day et al., 2020). PLA was performed on *3xflag::csn-5; gfp::fbf-1* and *3xflag::csn-5; gfp::fbf-2* animals using *3xflag::csn-5; gfp* as a control for spurious proximity. We observed a statistically significant (*p*<0.0001; Fig. 3) increase in PLA signal within the SPC zone in *3xflag::csn-5; gfp::fbf-2* compared to both control and *3xflag::csn-5; gfp::fbf-1.* Curiously, there was no significant difference in PLA signal between *3xflag::csn-5; gfp::fbf-1* and control. Although PLA signal with FBF-1 was not statistically different from background, it’s possible that FBF-1 could transiently interact with CSN-5, and this assay might not effectively capture those types of interactions. Nonetheless, this data suggests that FBF-2 interacts with CSN-5 *in vivo*.

**Fig. 3.**
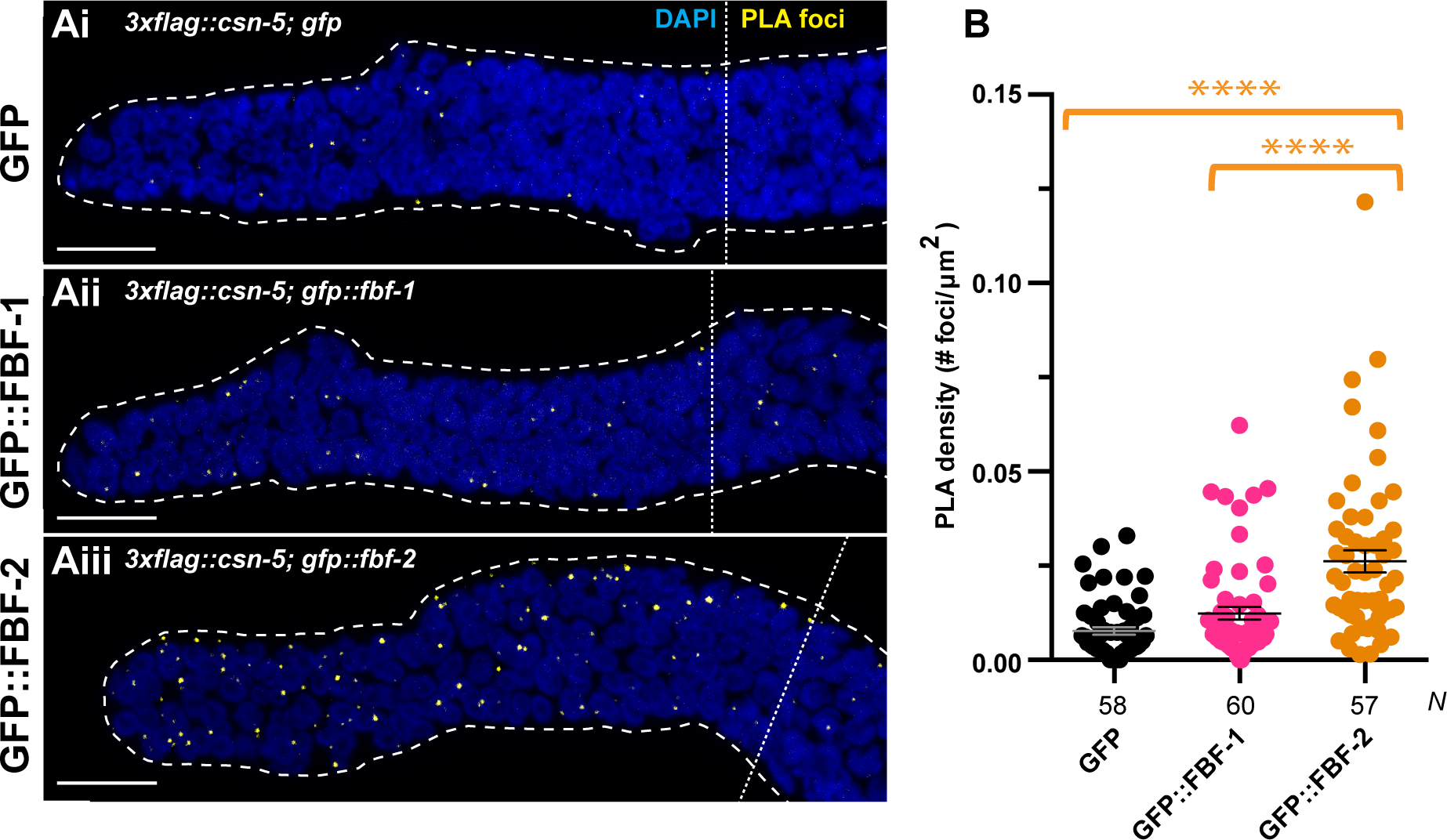
FBF-2 interacts with CSN-5 *in vivo*. (A) Confocal images of the distal germline SPC zones with PLA foci (yellow) and DAPI (blue). Germlines are outlined with dashed lines and vertical dotted lines indicate the beginning of the transition zone. Genotypes are indicated with their respective images. Scale bars: 10 µm. (B) The PLA density (number of PLA foci per µm^2^) within the SPC zone was measured for germlines of each genetic background. Differences in PLA density for each protein was evaluated by one-way ANOVA with Tukey’s post-test. Asterisks denote statistical significance (****, p<0.0001) where FBF-2 had significantly more PLA foci than both FBF-1 and the control. Number of germlines scored (*N*) are indicated below the graph. Data is representative of 3 biological replicates and error bars denote SEM. All experiments were performed at 24°C.

### COP9 is required to maintain FBF-1/2, but not PUF-3 protein levels in stem and progenitor cells

In *C. elegans*, CSN-5 was previously identified as an interacting partner of germline proteins, GLH-1 and GLH-3 (Smith et al., 2002), and found to promote accumulation of GLH-1 (Orsborn et al., 2007). Therefore, we hypothesized that CSN-5 might similarly promote FBF accumulation and aimed to test FBF levels in the germlines where *csn-5* function was disrupted. The effect of CSN-5 on FBFs may or may not depend on the COP9 complex (Tomoda et al., 1999; Wei et al., 2008; Yoshida et al., 2013). To distinguish COP9-dependent and independent roles of CSN-5, we additionally documented FBF levels in mutants of two COP9 subunits, *csn-2* and *csn-6*. If the whole complex were required for specific functions, we would expect to see the same, or very similar, phenotypes across all *csn(loss-of-function, lf)* mutants, as all mutations used in this study are expected to be null alleles (Brockway et al., 2014; Fig. S6; Materials and Methods). Furthermore, previous studies indicate that without CSN2 and CSN6, COP9 complex cannot readily incorporate CSN5, and thus remains inactive (Lykke-Anderson et al., 2003; Birol et al., 2014; Lingaraju et al., 2014). Additionally, it is unlikely that adult *csn(lf)* germlines would have residual CSN protein from maternal contribution (Oron et al., 2002). We quantified the total FBF protein levels in adult *csn(lf)* mutant worms by Western blotting with antibodies to the endogenous FBF-1 or epitope-tagged endogenous 3xV5::FBF-2 (Shin et al., 2017), and used tubulin as a loading control. We found the most drastic reduction of FBF-1 protein level in *csn-5(lf)* (0.24-fold of the wild type levels; *p*<0.005; Fig. 4A, B), whereas *csn-6(lf)* and *csn-2(lf)* were affected weaker (0.39-fold and 0.68-fold; *p*<0.01; *p*<0.05, respectively; Fig. 4A, B). Furthermore, FBF-1 protein levels in the *csn-5(lf)* mutant were significantly lower than those in the *csn-2(lf)* mutant (*p*<0.05; Fig. 4B). Immunostaining of the *csn-5(lf) and csn-6(lf)* mutants confirmed the loss of signal in the distal SPCs (Fig. 4D-G), where FBFs are normally expressed, and suggested that the reduction of FBF-1 protein does not simply result from a decrease in stem cell numbers. The stronger FBF-1 protein reduction in *csn-5(lf)* implies that CSN-5 might be promoting the accumulation of FBF-1 independently of the COP9 complex, since the relative amounts of FBF-1 would be similar across all mutants otherwise. Surprisingly and distinctly, we observed a significant reduction of 3xV5::FBF-2 in all *csn(lf)* mutants by Western blot (0.29-fold; *p*<0.0001; Fig. 4A, C) and confirmed this observation by immunostaining (Fig. 4H-K), suggesting COP9-dependence. We conclude that CSN-5, as well as COP9, promote FBF steady-state accumulation.

**Fig. 4.**
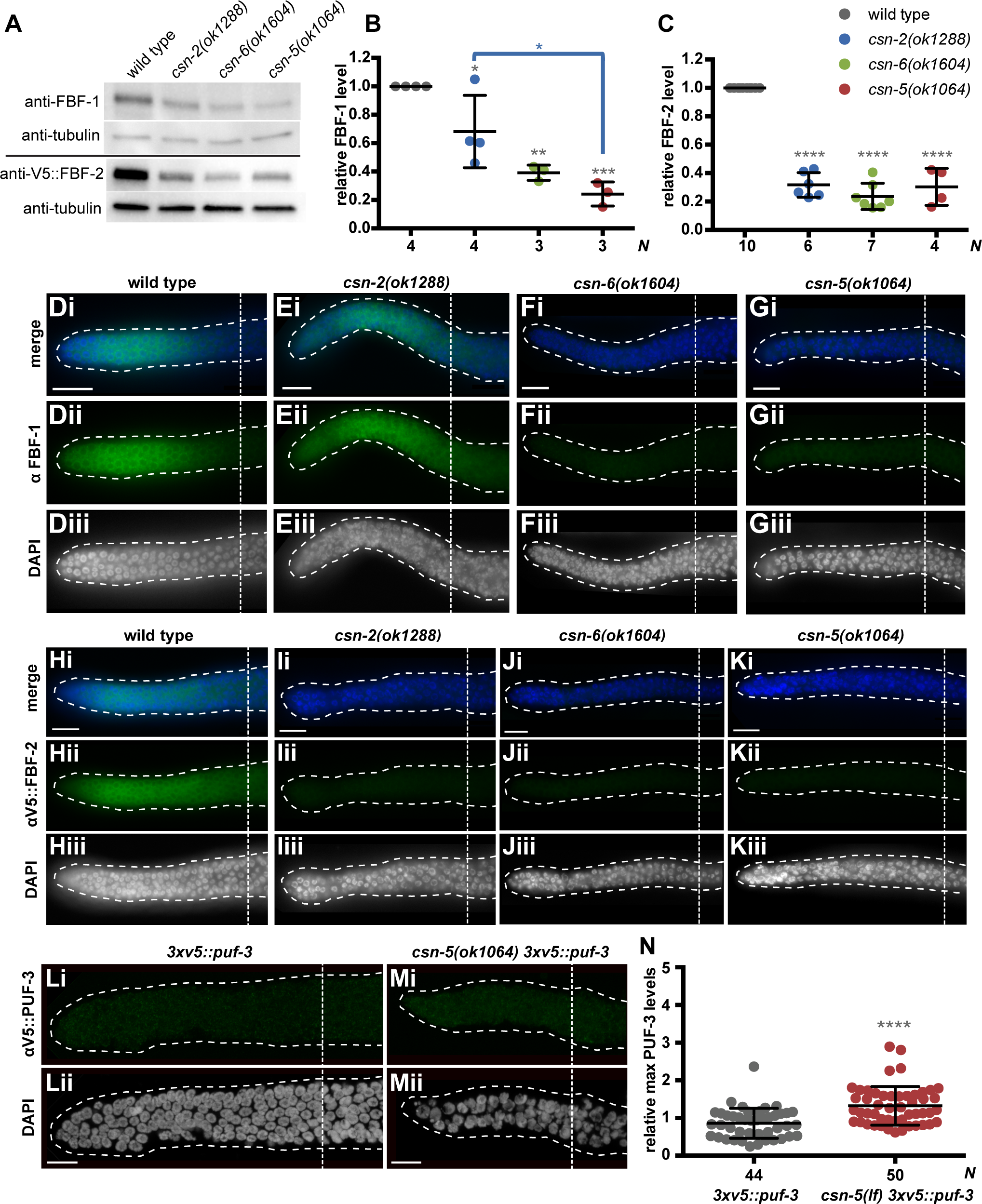
FBF-1/-2, but not PUF-3, protein levels are reduced in *csn(lf)* mutants. (A) Western blot analysis of *3xv5::fbf-2(q932),* referred to here as wild type, and respective *3xv5::fbf-2(q932); csn(lf)* mutant worm lysate reveals reduced levels of FBF-1 (top panels) and 3xV5::FBF-2 (bottom panels). Tubulin is used as a loading control. Endogenous FBF-1 and epitope-tagged endogenous 3xV5::FBF-2 are detected by anti FBF-1 and anti-V5. (B, C) Total FBF protein level of each *3xv5::fbf-2(q932); csn(lf)* mutant normalized to *3xv5::fbf-2(q932)*, referred to here as wild type. Differences in protein level were evaluated by one-way ANOVA with Tukey’s post-test. Grey asterisks denote statistical significance compared to wild type and blue bracket and asterisk denote statistical significance comparing *csn-2(lf)* to *csn-5(lf)* (*, *p*<0.05; **, *p*<0.01; ***, *p*<0.005; ****, *p*<0.0001). Number of biological replicates (*N*) is indicated at the bottom of the graphs. Mean group values are shown as lines and error bars denote standard deviation. (D-K) Gonads of *csn(lf)* mutants are dissected and stained with (D-G) anti FBF-1 (green) or (H-K) anti-V5 (green) and DAPI (blue) to visualize protein levels of FBF-1/-2. The individual FBF (green) and DAPI (greyscale) channels are also shown for better contrast. (L, M) Distal germlines dissected from adult *3xv5::puf-3* and *csn-5(lf) 3xv5::puf-3* dissected and stained with anti-V5 (green) and DAPI (grey). Germlines are outlined with dashed lines and vertical dotted lines indicate border of the SPC zone. Scale bars: 10 μm. (N) Quantification of SPC expression of 3xV5::PUF-3 protein. Maximum PUF-3 levels in each germline are scaled to average levels in the *3xv5::puf-3* strain. Differences in maximum protein level were evaluated by Student’s t-test. Grey asterisks denote statistical significance (****, *p*<0.0001). Number of germlines scored (*N*) is indicated at the bottom of the graphs. Mean group values are shown as lines and error bars denote standard deviation. Data reflect three biological replicates.

The reduction in FBF-1/2 protein levels may be caused by a decrease in *fbf* transcripts or by an effect on FBF proteins. To start distinguishing these possibilities, we tested if *csn(lf)* affects the steady-state levels of *fbf-1/-2* mRNAs using quantitative PCR. Using *unc-54* (myosin heavy chain) as a control and normalizing mRNA abundance to a reference gene, *act-1* (actin), we observed a reduction in *fbf-1* mRNA abundance in each of the mutants but due to high variability the difference was not statistically significant (Fig. S2). However, we observed a statistically significant reduction of *fbf-2* mRNA abundance in each *csn(lf)* mutant compared to wild type (*p*<0.0001; Fig. S2). By contrast, we did not observe a decrease in *unc-54* mRNA in any *csn(lf)* mutant (Fig. S2). As the effects of all *csn(lf)* mutants on the levels of *fbf* transcripts were similar, we conclude that COP9 signalosome is required for normal steady state levels of *fbf* mRNAs. However, distinct levels of FBF-1 protein in *csn-2(lf)* vs. *csn-5(lf)* mutant backgrounds (Fig. 4A, B) suggest that CSN-5 might additionally promote FBF-1 protein accumulation.

In addition to FBF-1 and FBF-2, *C. elegans* germline stem and progenitor cells express three other members of PUF-family RNA-binding proteins, PUF-3, PUF-11, and PUF-8 (Haupt et al., 2020; Ariz et al., 2009; Racher and Hansen, 2012). Although we were unable to determine if CSN-5 interacts with other PUF proteins *in vitro*, we aimed to test whether the levels of another PUF were affected by *csn-5(lf)*. We investigated PUF-3, for which a tagged *3xv5* allele is available (Haupt et al., 2020). PUF-3 is expressed at low levels in the progenitor zone and reaches its strongest expression in the oocytes (Haupt et al., 2020; Spike et al., 2022). For direct comparison with FBFs, we focused on the progenitor zone and immunostained *csn-5(lf) v5::puf-3* germlines with antibodies to V5-tag (Fig. 4L, M). We observed a significant increase in SPC 3xV5::PUF-3 levels (1.3 fold; *p*<0.0001; Fig. 4N) in the *csn-5(lf*) background, opposite of what we observe with FBF levels (Fig. 4B, C). We conclude CSN-5 contribution to protein accumulation is specific for FBFs in contrast to other PUF-family members, such as PUF-3.

### COP9 subunit mutants have reduced numbers of stem cells that continue to proliferate and enter meiosis

If CSN-5 facilitates FBF function in the germline, we expect *csn-5(lf)* mutation to cause developmental defects similar to those caused by *fbf-1/2(lf)*. Because FBFs regulate the switch from mitosis to meiosis in the *C. elegans* germline (Crittenden et al., 2002; Wickens et al., 2002) and therefore affect the size of its stem and progenitor cell (SPC) population, we quantified the number of SPCs in *csn(lf*) mutants. We dissected and stained adult germlines with the antibodies specific to proliferating cells in the progenitor zone, anti-REC-8, to visualize the SPC zone (Fig. 5Ai-iv; Hansen et al., 2004). We observed that all *csn(lf)* mutants had reduced numbers of SPCs compared to the wild type (*p*<0.0001, Fig. 5B), which is in general agreement with a previous report (Brockway et al., 2014). The *csn-6(lf)* and *csn-5(lf)* mutants had the most drastic reduction in SPCs, whereas the *csn-2(lf)* SPCs were less affected (*p*<0.01, Fig. 5B). Although a decrease in SPCs is not a phenotype unique to *fbf(lf)*, it is consistent with compromised FBF function. To test whether *csn-5* functions in the same genetic pathway as *fbfs*, we quantified the number of SPCs in *csn-5(lf)* following *fbf* knockdown with RNAi that targets both *fbf-1* and *fbf-2* (we were not able to generate a triple-mutant strain because of incompatibility between *mIn1* and *nT1* balancers). When cultured at higher temperatures (24°C), *fbf(RNAi)*-treated animals maintain reduced numbers of SPCs (Merritt and Seydoux 2010; Hansen and Schedl 2013). Efficacy of RNAi treatment was confirmed by germline masculinization of *csn-5(lf)/nT1* as seen in *fbf-1/-2(lf)* mutants (Crittenden et al., 2002). We observed no additive effect on SPC number in *csn-5(lf) fbf(RNAi)* background (Fig. S3A), supporting the hypothesis that *csn-5* functions in the same genetic pathway as *fbf-1* and *fbf-2* in germline SPCs.

**Fig. 5.**
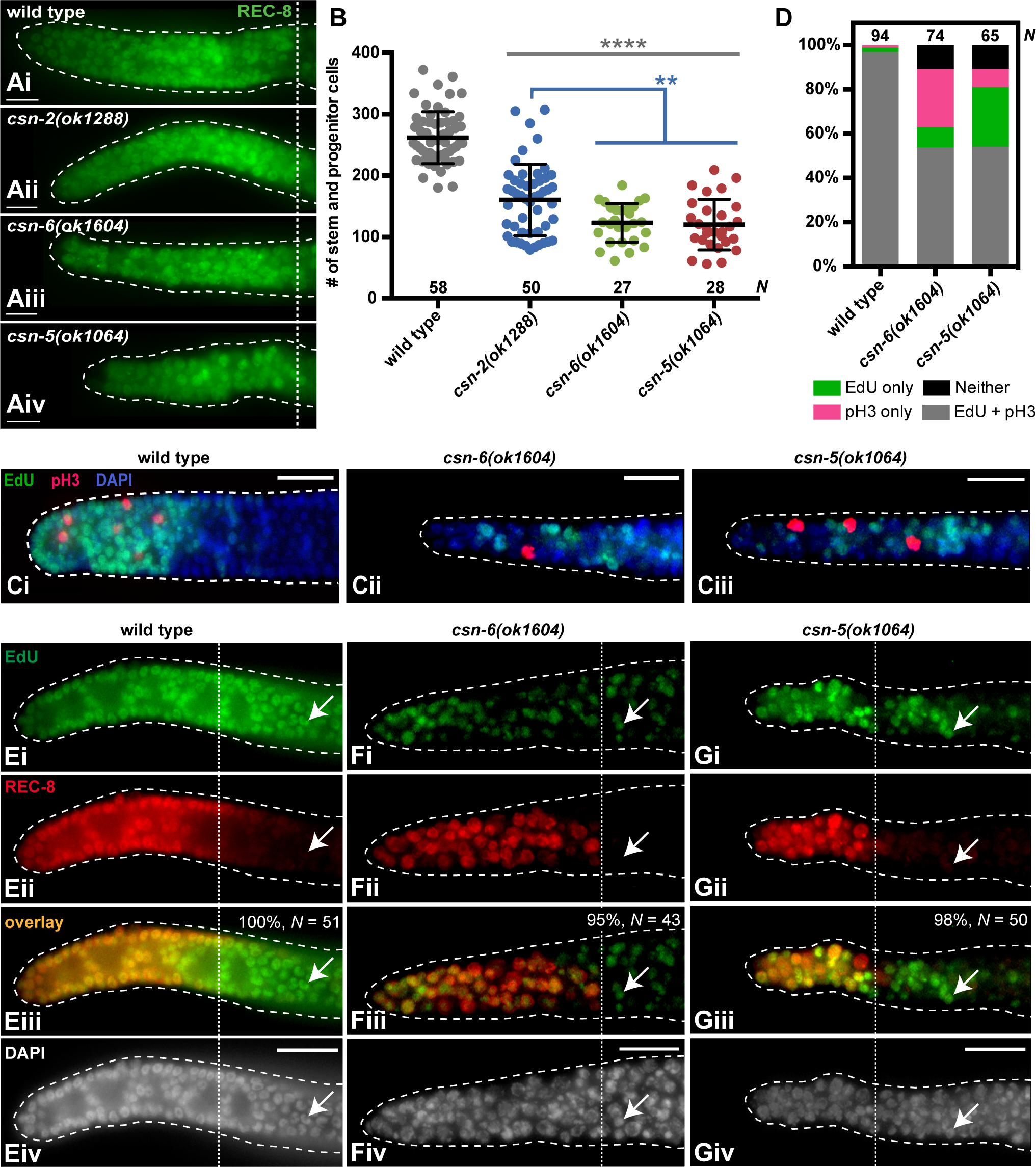
COP9 subunit mutants have reduced number of stem cells but sustain proliferation and meiotic entry. (Ai-iv) Distal germlines dissected from adult *3xv5::fbf-2(q932)*, referred to as wild type, and respective *3xv5::fbf-2(q932); csn(lf)* mutants were stained with anti-REC-8 (green) to visualize SPC zone. Germlines are outlined with dashed lines and vertical dotted lines indicate the SPC zone boundary. Scale bars: 10 μm. (B) Quantification of SPCs in wild type and *csn(lf)* mutants. Differences in number of SPCs was evaluated by one way ANOVA with Tukey’s post-test. Grey asterisks denote statistical significance compared to wild type (****, *p*<0.0001) and blue asterisks denote statistical significance compared to *csn-2(lf)* (**, *p*<0.01). Number of germlines scored (*N*) are indicated at the bottom of the graph. Stem cell quantification reflects six biological replicates of wild type and three biological replicates for each *csn(lf)* mutant. (Ci-iii) Distal germlines dissected from adult *3xv5::fbf-2(q932)*, referred to as wild type, and respective *3xv5::fbf-2(q932); csn(lf)* mutants were stained for EdU and pH3 to visualize cells in S-phase and identify mitotically dividing cells, respectively. Germlines are outlined with dashed lines. Scale bars: 10 μm. (D) Germlines were scored for the presence of EdU incorporation, pH3(+) cells, both, or neither. Number of germlines scored (*N*) are indicated at the top of each column. Data are plotted in aggregate and reflect two biological replicates. (E-G) Distal germlines from adult *3xv5::fbf-2(q932)*, referred to as wild type, and respective *3xv5::fbf-2(q932); csn(lf)* mutants dissected and stained for EdU and REC-8 to visualize cells in S-phase and the SPC zone, respectively. Arrows indicate EdU(+)/REC-8(-) cells entering meiosis. Corresponding percent of germlines entering meiosis with number of germlines scored (*N*) indicated in overlay panels (iii), data reflect two biological replicates. Germlines are outlined with dashed lines and vertical dotted lines indicate the SPC boundary. Scale bars: 10 μm.

Because *csn-5(lf)* and *csn-6(lf)* mutant germlines are visibly smaller than wild type (Fig. 5Aiii-iv, and full germline images in Fig. 6Aiii-iv), we tested whether their germline SPCs arrested cell cycle progression. To test whether germline SPCs continue to proliferate and enter meiosis, we performed EdU incorporation assays combined with immunostaining with antibodies specific for mitotic M-phase or proliferating cells in the progenitor zone. First, young adult germlines were labeled with the thymidine analog 5-ethinyl-2’-deoxyuridine (EdU) for 4 hours and stained for phospho-histone H3 (pH3) to identify cells in S-phase and M-phase respectively, and scored by how many germlines were positive for: pH3, EdU incorporation, both pH3 and EdU incorporation, or had neither (Fig. 5C, D). We found that approximately 90% of *csn-5/-6(lf)* germlines had SPCs positive for EdU incorporation and/or pH3, together, indicating active cell cycle progression. Next, we tested if *csn-5/-6(lf)* SPCs were capable of meiotic entry by staining adult gonads that have been labeled with EdU for 14 hours with REC-8, and scoring how many germlines had EdU-positive cells that were able to initiate meiosis and move beyond the SPC zone as detected by the loss of REC-8 staining (Fox et al., 2011; Kocsisova et al., 2018). Approximately 98% and 95% of *csn-5/-6(lf)* germlines, respectively, exhibited progression of EdU-positive cells into meiosis (Fig. 5E-G). Additionally, we observed weaker EdU signal in the two *csn(lf)* mutants and found that EdU-positive cells accumulated slower than in the wild type despite identical EdU treatment conditions (data not shown, see Materials and Methods for detail). We conclude that *csn-5(lf)* and *csn-6(lf)* germline SPCs sustain proliferation and meiotic entry, although both cell cycle and meiotic entry appear slower than in the wild type.

### Defective oogenesis in COP9 subunit mutants

FBFs play a role in germline sex determination by regulating the spermatogenesis to oogenesis switch (Zhang et al., 1997; Crittenden et al., 2002), so we assessed if germline sex determination was affected in the *csn(lf)* mutants. The majority of *csn-6(lf)* and *csn-5(lf)* mutants failed to produce oocytes and generated only sperm (Fig. 6Aiii-iv, B). This raised the possibility that downregulation of FBF protein in these *csn(lf)* genetic backgrounds might be secondary to germline masculinization, as previously observed for GLD-1 (Jones et al., 1996). To address this, we compared FBF levels in the male versus hermaphrodites of the same genetic background and observed no significant difference in FBFs (Fig. S4A), suggesting that the decrease in FBF protein levels is likely a direct consequence of the *csn-5(lf)* and *csn-6(lf)*. Although germline masculinization can result from a variety of genetic causes (reviewed in Ellis, 2021), it is consistent with a decrease in *fbf* function (Zhang et al., 1997; Crittenden et al., 2002). Curiously, the *csn-2(lf)* mutant had a clearly distinct phenotype where most germlines could still form oocytes (Fig. 6Aii, B). As we observed significantly more FBF-1 protein in *csn-2(lf)* than in *csn-5(lf)* (Fig. 4B), we next combined *csn-2(lf)* with *fbf-1(lf)* to test if the switch to a female germ cell fate in *csn-2(lf)* was dependent on FBF-1. We found the levels of masculinization in *csn-2(lf); fbf-1(lf)* germlines increased to 82%, similar to those observed in *csn-5(lf)* (83%; Fig. 6B). This is consistent with the interpretation that the reduction of FBFs underlies disrupted sperm/oocyte switch in *csn-5/-6(lf*) mutants.

**Fig. 6.**
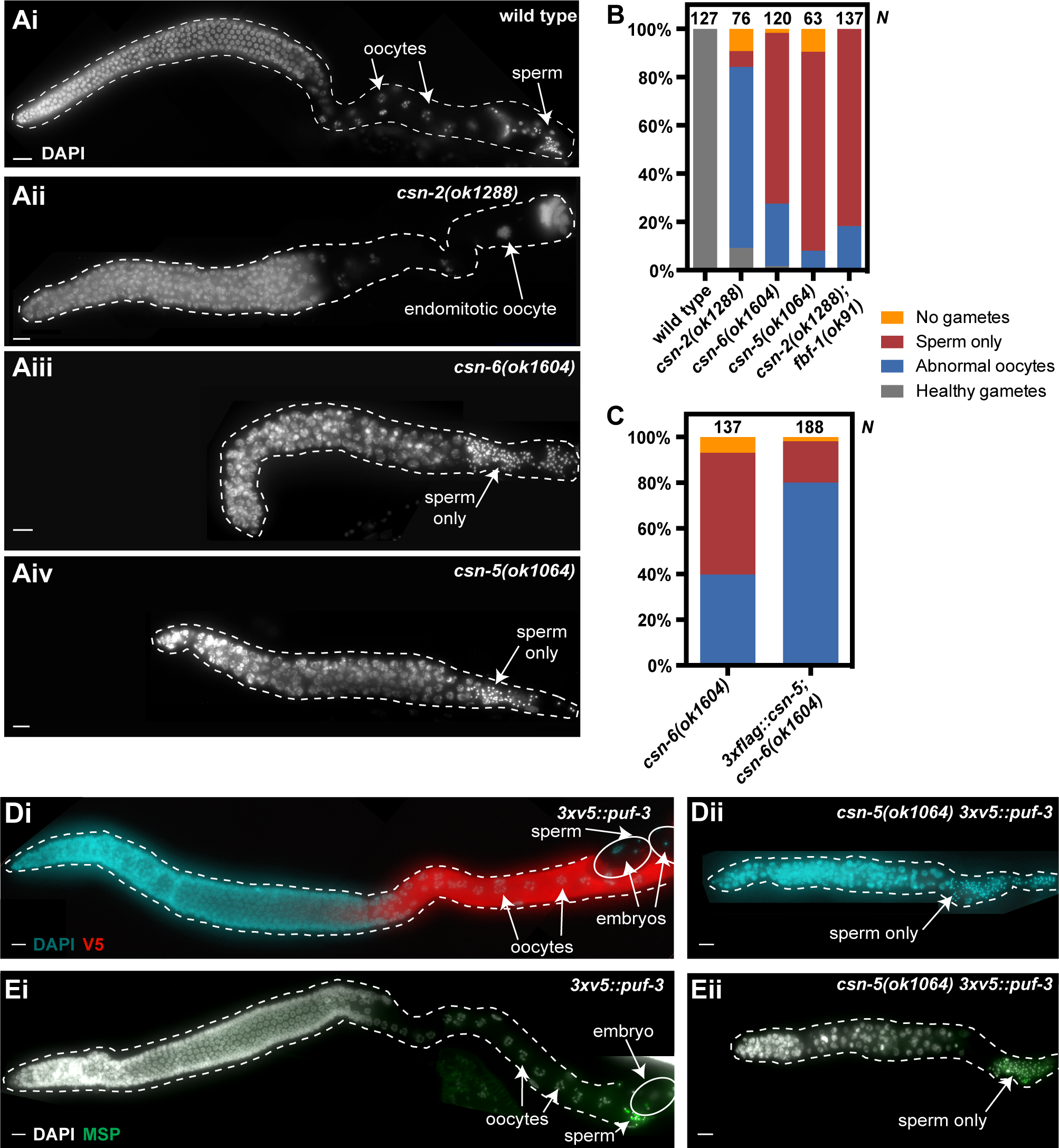
Mutations in COP9 subunits disrupt oogenesis. (Ai-iv) Gonads of adult *3xv5::fbf-2(q932)*, referred to as wild type, and respective *3xv5::fbf-2(q932); csn(lf)* mutants indicated in each panel dissected and stained for DNA with DAPI. Arrows indicate gametes. Germlines are outlined with dashed lines. Scale bars: 10 μm. (B) Gametogenesis in wild type and *csn(lf)* mutants. Number of germlines scored (*N*) are indicated at the top of each column. Data are plotted in aggregate and reflect six biological replicates of wild type, four replicates of *csn-6(lf)*, three replicates of each *csn-2(lf)* and *csn-5(lf)*, and two replicates of *csn-2(lf); fbf-1(lf)*. (C) Female fate of *csn-6(lf)* is partially restored by an extra copy of *csn-5* gene in *3xflag::csn-5; csn-6(lf)* strain. Number of germlines scored (*N*) are indicated at the top of the graph. Data are plotted in aggregate and reflect three biological replicates. (D, E) Gonads of adult *3xv5::puf-3* and *csn-5(lf) 3xv5::puf-3* mutant indicated in each panel dissected and stained for DNA with DAPI (cyan/grey), V5::PUF-3 with anti-V5 (D, red), or sperm with anti-MSP (E, green). Arrows indicate gametes. Germlines are outlined with dashed lines. Scale bars: 10 μm.

To investigate molecular markers of oogenesis in *csn-5(lf)* germlines, we examined expression of PUF-3 that is highly expressed in proximal pachytene at the start of oogenesis and throughout oocytes (Haupt et al., 2020; Spike et al., 2022; Fig. 6Di). We observed the absence of 3xV5::PUF-3 at the proximal end of *csn-5(lf)* germlines in the presence of sperm, suggesting a failure of the switch to oogenesis (Fig. 6Dii, Eii). This supports our initial interpretation that *csn-5(lf)* mutant germlines fail to promote female fate, rather than arrest oogenic pachytene cells at the transition to diplotene.

### CSN-5’s role in germline sex determination is largely independent of COP9 and cell autonomous

We hypothesized that the similar germline stem cell and sex determination phenotypes observed in *csn-5(lf)* and *csn-6(lf)* mutants were caused by a decrease in the levels of CSN-5 protein in *csn-6(lf)*. Indeed, we observed that the levels of 3xFLAG::CSN-5 transgene in a *csn-6(lf)* background decrease by approximately 60% compared to the control (Fig. S4B), similar to a previous observation of a decrease in endogenous CSN-5 levels after a knockdown of *csn-6* (Miller et al., 2009). We aimed to test whether CSN-5-dependent, COP9-independent phenotypes might be rescued by increasing *csn-5* copy number in the *csn-6(lf)* background. To this end, we first tested whether the *3xflag::csn-5* transgene was able to complement the *csn-5(lf)* mutant and observed complete rescue of fertility (Table 1). Next, we introduced the *3xflag::csn-5* transgene into *csn-6(lf)* and found that an extra copy of *csn-5* was able to partially rescue germline masculinization of *csn-6(lf)*, resulting in phenotypes similar to those of *csn-2(lf)* (Fig. 6C). However, it was unable to rescue the low numbers of stem cells in *csn-6(lf)* (data not shown). Ectopic CSN-5 in *csn-6(lf)* would be unable to rescue COP9 function because in the absence of CSN-6, CSN-5 fails to incorporate into the COP9 holoenzyme and the complex remains enzymatically inactive (Birol et al., 2014; Lingaraju et al., 2014). We conclude that the role of CSN-5 in FBF-mediated sex determination does not require COP9 signalosome.

**Table 1:** Germline-expressed CSN-5 rescues fertility defect of *csn-5(lf)*

| Genotype | % Fertile | *n* |
| --- | --- | --- |
| *3xv5::fbf-2(q932); csn-5(ok1064)* | 0 | >500 |
| *csn-5p::3xflag::csn-5; csn-5(ok1064)* | 100 | >500 |
| *gld-1p::3xflag::csn-5; csn-5(ok1064)* | 100 | 311 |
| *gld-1p::3xflag::csn-5(D152N); csn-5(ok1064)* | 46.4* | 110 |

*n* indicates number of germlines scored.
\* Fertile animals were able to produce embryos that failed to hatch

Because CSN-5 is expressed in both germline and somatic tissues (Smith et al., 2002; Miller et al., 2009), we aimed to test whether its function was required in the soma or the germline. We generated a *3xflag::csn-5* transgene using the *gld-1* promoter, which is expected to restrict expression to the germline (Ellenbecker et al., 2019; see Materials and Methods for details), and still observed 100% rescue of fertility when incorporated into the *csn-5(lf)* mutant background (Table 1). This suggests that the effects of *csn-5* on sex determination are likely cell-autonomous within the germline. We next tested whether the *csn-5* function in promoting female germ cell fate was dependent on its deneddylating activity by generating a catalytically inactive CSN-5(D152N) transgene, based on a previously reported catalytic mutant (Cope et al., 2002; Peth et al., 2007). CSN-5(D152N) is incorporated into COP9, but fails to promote de-neddylation (Cope et al., 2002); therefore, it would only rescue COP9-independent function of *csn-5*. We find that 3xFLAG::CSN-5(D152N) transgene can partially rescue oogenesis in *csn-5(lf)* (Table 1), suggesting that the effect of *csn-5* on germline sex determination is not entirely dependent on its proteolytic activity. However, since CSN-5-mediated deneddylation by COP9 is required for embryonic cell division (Pintard et al., 2003), all embryos produced by the *3xflag::csn-5(D152N); csn-5(lf)* were dead. Additionally, inactive CSN-5(D152N) maintains its ability to bind FBF-2 *in vitro* via GST pulldown (Fig. S4C) as expected if its effect depended on interaction with FBF-2.

### CSN5 maintains FBF protein levels through multiple mechanisms

To begin investigating a mechanism behind the compromised FBF accumulation and function in *csn-5(lf)* mutants, we tested whether CSN-5 stabilizes FBF proteins by interfering with their proteasome-mediated degradation. If true, we expect that disruption of the proteasome function in *csn-5(lf)* mutants would restore FBF protein levels. To test this hypothesis, we knocked down *pbs-6*, a catalytic subunit of the 20S proteasome, in the *rrf-1(lf)* genetic background to preferentially direct the RNAi knockdown to the germline (Sijen et al., 2001; Kumsta and Hansen, 2012). We observed a complete rescue of both steady-state FBF-1 (*p*<0.01) and 3xV5::FBF-2 (*p*<0.05) protein levels in *rrf-1(lf); csn-5(lf)* following *pbs-6(RNAi)* (Fig. 7A-D). We were unable to assess the effects of this rescue on SPC zone size as *pbs-6(RNAi)* on the control strain *rrf-1(lf); 3xv5::fbf-2(q932)* reduced the number of SPCs similar to *csn-5(lf)* (data not shown). Unexpectedly, crossing *csn-5(lf)* into *rrf-1(pk1417)* genetic background largely suppressed the failure of oogenesis and allowed formation of abnormal oocytes in 79% of examined germlines precluding testing whether *pbs-6(RNAi)* rescued oogenesis (Fig. S5). Understanding the nature of this suppression, including whether it results from the loss of *rrf-1* or a different background or linked mutation, remains the subject of future research. Since we previously observed a decrease in *fbf-1/2* transcript levels in *csn-5(lf)* background (Fig. S2), we next determined whether the rescue of FBF protein accumulation upon proteasome disruption was mediated by an increase in *fbf* mRNA abundance. Quantification of steady-state *fbf* mRNA levels by qRT-PCR revealed that *pbs-6(RNAi)* did not increase *fbf* transcript levels (Fig. 7E). We conclude that CSN-5 promotes the stability of FBF proteins.

**Fig. 7.**
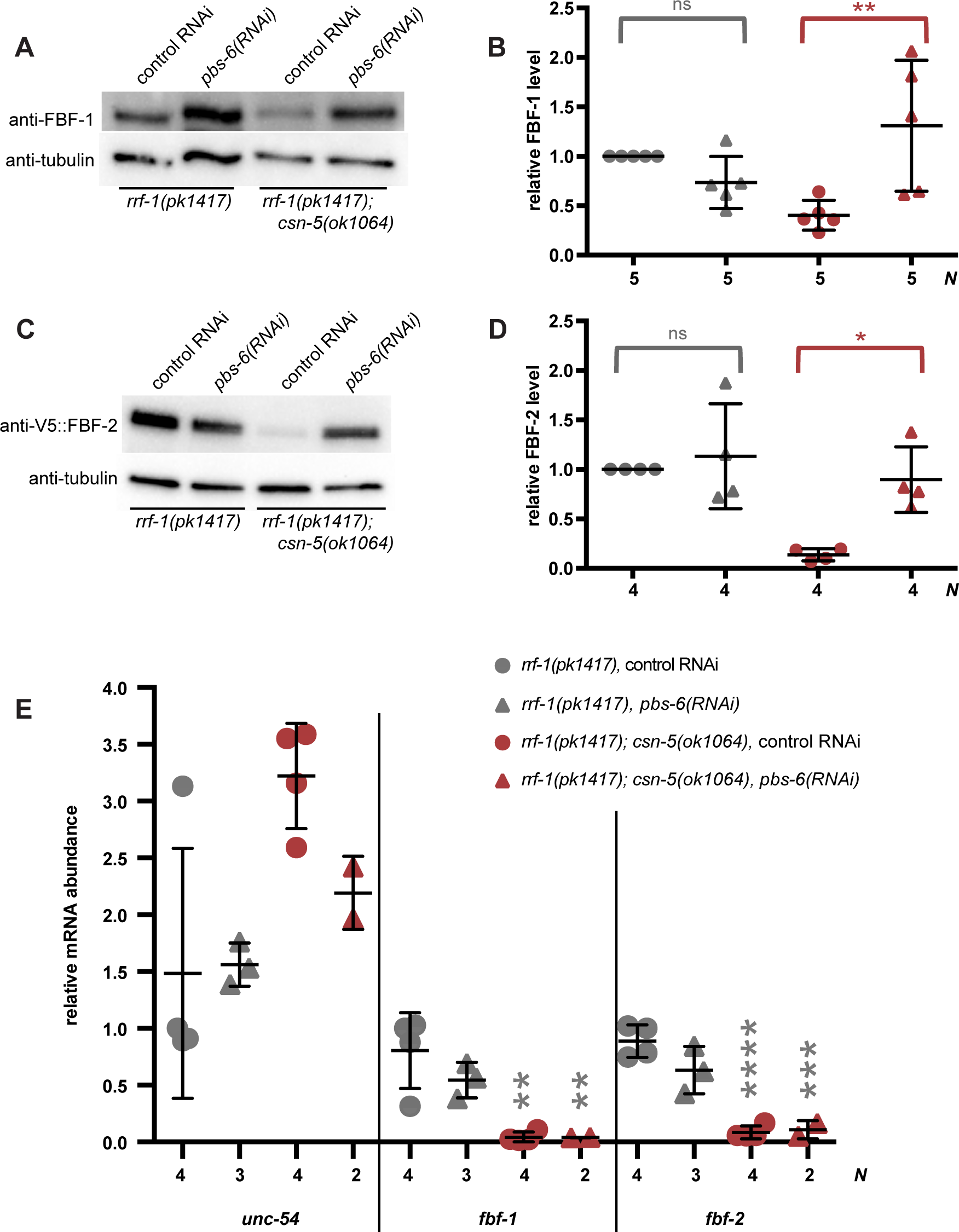
*csn-5(lf)* destabilizes FBF proteins. (A, C) FBF protein levels after treatment with a control RNAi and *pbs-6(RNAi)* in *rrf-1(lf); 3xv5::fbf-2(q932)*, referred to as *rrf-1(pk1417)* here, and *rrf-1(lf); 3xv5::fbf-2(q932); csn-5(lf)*, referred to as *rrf-1(pk1417); csn-5(ok1064)*. (A) Endogenous FBF-1 and (C) epitope-tagged endogenous 3xV5::FBF-2 are detected by anti-FBF-1 and anti-V5. Tubulin is used as a loading control. (B, D) Total FBF protein levels of *csn-5(lf)* normalized to wild type control RNAi. Control RNAi data points for each genetic background are shown as circles and *pbs-6(RNAi)* is shown as the triangles. Difference in FBF protein level was evaluated by one-way ANOVA with Sidak’s post-test. Red brackets with asterisks denote statistical significance comparing *csn-5(lf)* control RNAi to *csn-5(lf) pbs-6(RNAi)* (*, *p*<0.05; **, *p*<0.01), ns denotes not significant. Number of biological replicates (*N*) is indicated below the graph, mean value is indicated as a line, and error bars denote standard deviation. (E) Steady-state levels of transcripts indicated on the X-axis are quantified by RT-qPCR and normalized to reference gene *act-1*; *unc-54* is a control. Differences in relative mRNA abundance of *fbf-1/-2* were evaluated by one-way ANOVA with Dunnett’s post-test. Asterisks denote statistical significance compared to wild type control RNAi (**, *p*<0.01; ***, *p*<0.005; ****, *p*<0.0001). The data reflect four biological replicates of control RNAi for *rrf-1(lf)* and *rrf-1(lf); csn-5(ok1064),* three replicates of *rrf-1(lf); pbs-6(RNAi)*, and two replicates of *rrf-1(lf); csn-5(ok1064); pbs-6(RNAi)*. Mean values are plotted as lines with error bars representing standard deviation.

The main enzymatic activity of the COP9 signalosome is deneddylation of cullin subunits of **S**kp1-**c**ullin 1-**F**-box (SCF) ubiquitin ligases to promote turnover of the SCF targets (Lyapina et al., 2001; Cope et al., 2002; Pintard et al., 2003). Therefore, some phenotypes of *csn(lf)* mutants could result from the increased neddylation. Notably, a knockdown of *C. elegans* NEDD8 homolog, *ned-8*, has been shown to partially suppress meiotic defects seen in *csn(lf)* mutants (Brockway et al., 2014). By contrast, the effects of mammalian CSN5 on stability and function of its protein partners HIF-1a and E2F1 are independent of its deneddylase activity (Bemis et al., 2004; Hallstrom and Nevins, 2006). To investigate whether *csn-5* is affecting FBFs stability via deneddylation, we performed *ned-8(RNAi)* on the *csn-5(lf)* mutant (we were unable to generate a *ned-8; csn-5* mutant strain for analysis due to larval arrest observed in the homozygous *ned-8* mutant, Brockway et al., 2014; data not shown). We quantified expression of FBF-1 and V5::FBF-2 in distal mitotic cells simultaneously by co-staining the treated germlines with respective antibodies. Interestingly, we observed a partial rescue of FBF-2, but not FBF-1, levels in *ned-8(RNAi)*-treated *csn-5(lf)* mutants (increased to 0.76-fold on average; Fig. 8A-F). We conclude that CSN-5 contributes to FBF-1 and FBF-2 maintenance through two mechanisms, stabilizing FBF-1 independent of the deneddylating activity, and promoting FBF-2 accumulation through a combination of deneddylation-dependent and -independent mechanisms. Consistent with the partial rescue of FBF-2 function, we find that *ned-8(RNAi)* treatment of *csn-5(lf)* partially rescues SPCs numbers (Fig. 8G). Additionally, we observed that *ned-8(RNAi)* completely rescues oogenesis (Fig 8H). However, interpretation of this result is complicated by the fact that neddylation is essential for the function of cullin-based E3 ubiquitin ligases (Duda et al., 2008; Saha and Deshaies, 2008; Boh et al., 2011), and a cullin-based CUL-2/FEM-1/FEM-2/FEM-3 ubiquitin ligase is required for spermatogenesis (Starostina et al., 2007). Therefore, it is unclear whether the rescue of oogenesis is solely due to an increase in FBF-2 or also to downregulation of CUL-2/FEM-1/FEM-2/FEM-3 activity.

**Fig. 8.**
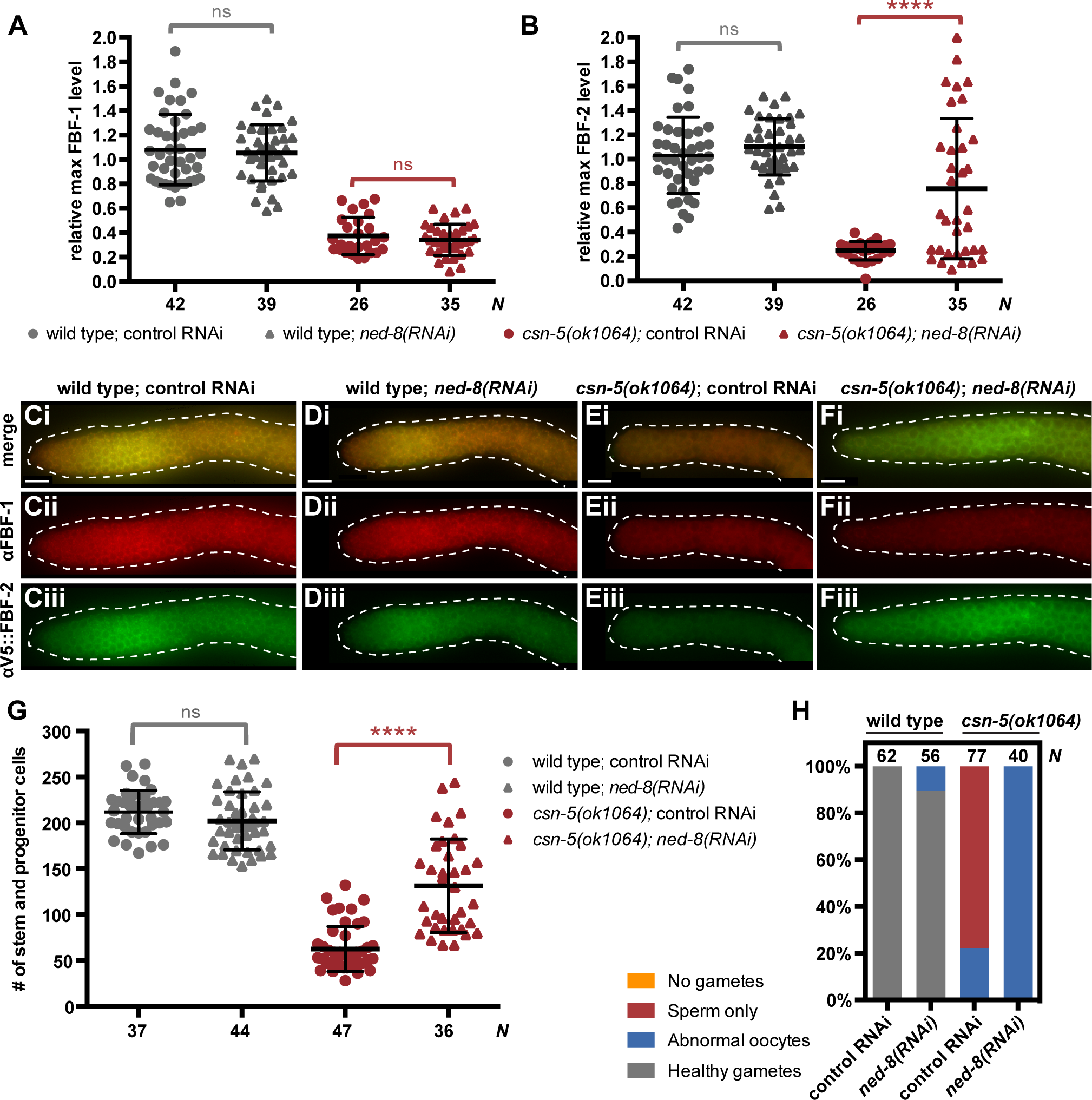
*ned-8(RNAi)* rescues phenotypes seen in *csn-5(lf)*. (A, B) Maximum intensity of FBF-1 (A) and FBF-2 (B) immunostaining of *3xv5::fbf-2(q932)*, referred to here as wild type, and *3xv5::fbf-2; csn-5(lf)*, referred to here as *csn-5(lf)* treated with control or *ned-8(RNAi)*. Endogenous FBF-1 and epitope-tagged endogenous 3xV5::FBF-2 detected by anti-FBF-1 and anti-V5. Maximal FBF protein intensity in each germline was normalized to the average of wild type control RNAi. Differences in protein level were evaluated by one-way ANOVA with Sidak’s post-test, ns denotes not significant, red asterisks denote statistical significance (****, *p*<0.0001). Number of germlines scored (*N*) is indicated at the bottom of the graphs. Mean group values are shown as lines and error bars denote standard deviation. The data reflect three biological replicates. (C-F) Gonads of wild type and *csn-5(lf)* mutant are dissected and stained with anti-FBF-1 (red) or anti-V5 (green) and DAPI (not shown) to visualize protein levels of FBF-1/-2. (G) Quantification of SPCs in wild type and *csn(lf)* mutants with *ned-8(RNAi)*. Differences in number of SPCs were evaluated by one-way ANOVA with Sidak’s post-test. ns denotes not significant and red asterisks denote statistical significance (****, *p*<0.0001). Number of germlines scored (*N*) are indicated at the bottom of the graph. Stem cell quantification reflects four biological replicates of *csn-5(lf)* control RNAi and three biological replicates of wild type control RNAi, wild type *ned-8(RNAi)*, and *csn-5(lf) ned-8(RNAi*). (H) Gametogenesis in *ned-8(RNAi)* treated wild type and *csn-5(lf)* mutant. Number of germlines scored (*N*) are indicated at the top of each column. Data are plotted in aggregate and reflect three biological replicates.

## Discussion

In this study, we have identified the COP9 signalosome component CSN-5 as a new interacting partner of PUF-family RNA-binding proteins FBF-1 and -2, and documented both COP9-independent and -dependent CSN-5 functions in promoting stem and progenitor cell (SPC) proliferation as well as female fate in *C. elegans* germline. Analysis of *csn-5(lf)* mutant background uncovered instability of FBF-1 and -2 proteins that is counteracted by CSN-5. We propose that CSN-5 maintains steady-state levels of FBF-1 and FBF-2 in the *C. elegans* germline by distinct mechanisms (Fig. 9).

**Fig. 9.**
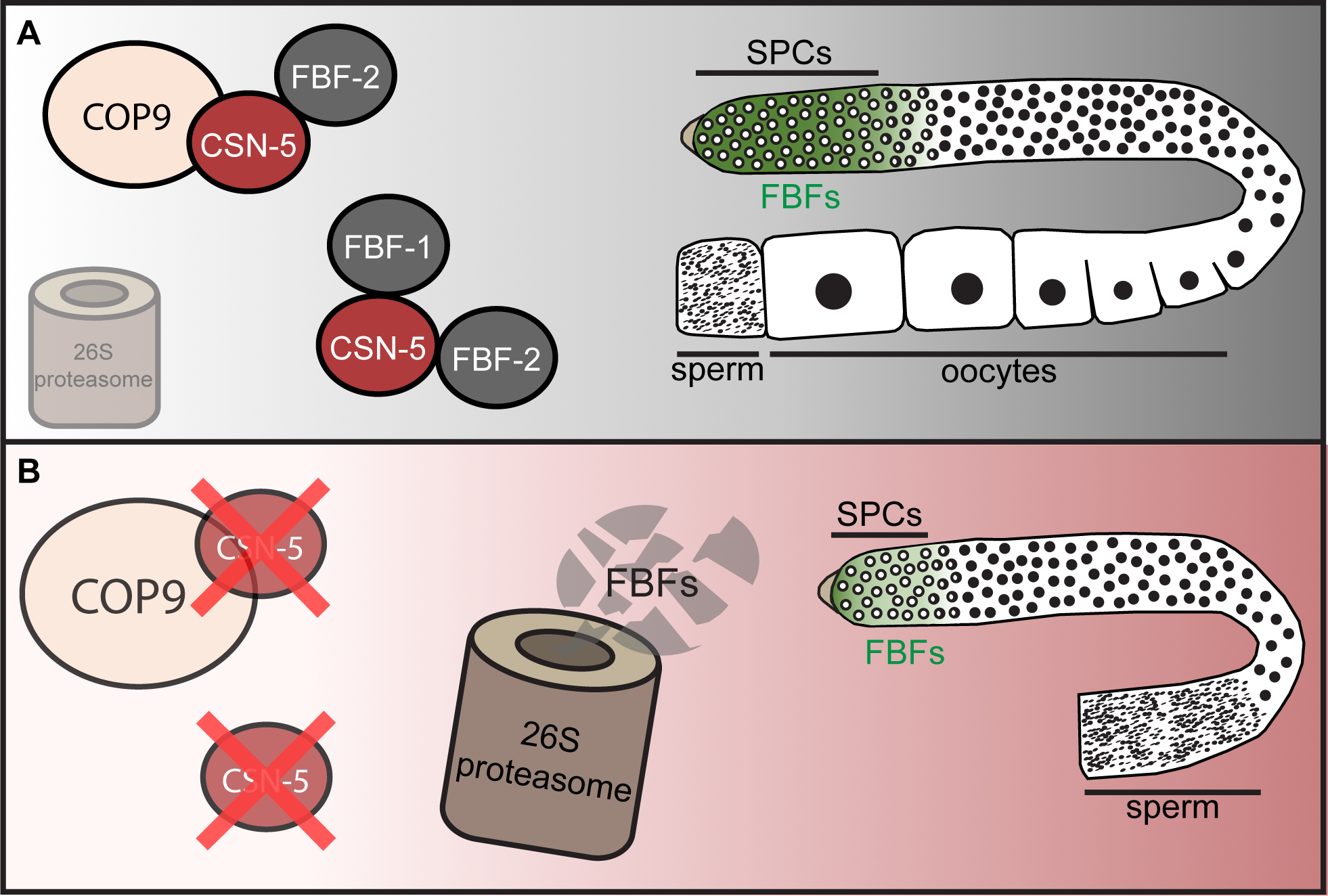
Model of CSN-5 stabilizing FBFs. (A) CSN-5 promotes the stability of FBFs both in the context of COP9 signalosome and independent of COP9, thereby allowing FBFs proteins to maintain the SPC zone and enable the switch from spermatogenesis to oogenesis. (B) Intrinsically unstable FBF proteins are readily degraded by the proteasome resulting in a germline with fewer SPCs and no female gametes in the absence of CSN-5.

### An interaction between MPN and PUF domains

We found the interaction between CSN-5 and FBFs in a yeast two-hybrid screen and confirmed it with GST pulldowns of recombinant proteins (Fig. 1B, D). Curiously, CSN-5 hasn’t been reported in the prior published yeast two hybrid screen using FBF-1 as a bait (Kraemer et al., 1999; Eckmann et al., 2002), but it might be a result of differences in prey library construction. We found that the direct interaction between PUF-family proteins (FBF-1, -2, and PUM1) and CSN5 homologs is mediated by conserved structured domains, the RBD and the MPN domain respectively (Fig. 2A, C, F). However, not all PUF RBDs bind CSN5 and not every type of MPN domain binds PUF proteins. At this time, the determinants of PUF/MPN binding are enigmatic. As reviewed in Wickens et al., 2002, PUF domains of PUM1 and PUM2 are very similar to each other (over 90% identical; Spassov and Jurecic, 2002), yet they exhibit distinct interaction properties with the CSN5^MPN^ (Fig. 2F). On the other hand, FBF-1 and FBF-2 belong to a separate, nematode-specific clade of PUF RBDs, with less than 30% identity to PUM RBDs (Wickens et al., 2002; Spassov and Jurecic, 2003), yet both are able to interact with CSN-5^MPN^. This raises the question whether additional PUF proteins out of ten described in *C. elegans* also bind to CSN-5. In addition to FBFs, germline stem cells express PUF-3 and PUF-11 that cluster into a nematode-specific subfamily related to FBFs (Haupt et al., 2020) and PUF-8 that is closer to human and *Drosophila* homologs (Ariz et al., 2009; Racher and Hansen, 2012). We were not able to test association between CSN-5 and PUF domains of PUF-3, -11, and -8 due to difficulties with recombinant PUF expression. If these interactions do exist, it is possible that some of the effects of *csn-5(lf)* are due to downregulation of additional germline PUFs. However, we do not observe a downregulation of PUF-3 in *csn-5(lf)* stem cells (Fig. 4L-N), suggesting specificity towards FBFs.

Similar to CSN-5, CSN-6 contains an MPN domain (Birol et al., 2014). However, CSN-6’s MPN domain could only bind FBF-2’s RBD at much higher concentrations than CSN-5 (Fig. 2E). Taken together, this suggests a selective association between a subset of PUF RNA-binding proteins and their specific MPN domain-containing partners. In the future, these selective interactions can help elucidate the sites mediating binding between these proteins.

We further find *in vivo* association between CSN-5 and FBF-2 by proximity ligation assay (Fig. 3). Although the *in vivo* proximity signal between CSN-5 and FBF-1 did not reach statistical significance, it’s possible that the interaction of CSN-5 with FBF-1 might be unstable or transient, and thus remain undetectable with PLA. In line with this, co immunoprecipitation has not recovered CSN-5 as an FBF-1 interactor (Friend et al., 2012). Additionally, both CSN-5 and FBF-2 are present in P granules and thus might interact more frequently compared to FBF-1, which does not localize to P granules (Smith et al., 2002; Voronina et al., 2012; Marnik et al., 2019). Alternatively, FBF-1 might be indirectly regulated by CSN-5. Nonetheless, here, we demonstrate that both FBFs require CSN-5 for proper accumulation and function.

### CSN-5 promotes FBF accumulation

CSN-5 and its homologs promote stabilization of their interaction partners (Bemis et al., 2004; Orsborn et al., 2007). We demonstrate that CSN-5 is similarly involved in stabilizing FBFs, as both FBFs are significantly reduced in *csn(lf)* mutants (Fig. 4A-K). We find that FBF-1 is most significantly reduced in *csn-5(lf)*, compared to *csn-6(lf)* or *csn-2(lf)* (Fig. 4A, B), suggesting that *csn-5* employs an additional mechanism promoting FBF-1 accumulation, independent of the COP9 complex. Conversely, we find that FBF-2 is depleted similarly in the *csn(lf)* mutants (Fig. 4A, C), suggesting the entire complex might promote FBF-2 protein accumulation. Interestingly, we also observed a reduction of *fbf* mRNAs in the *csn(lf)* mutants (Fig. S2), indicating that the entire COP9 complex is needed to maintain steady-state transcript levels. As reviewed in Chamovitz, 2009, a clear understanding of the COP9 complex in regulating transcription, outside the realm of stabilizing transcription factors and thus affecting steady-state transcript levels, remains somewhat elusive. Taken together, we conclude that there are both COP9-independent and –dependent functions of CSN-5 regulating FBF protein levels. We further demonstrate that the reduction of FBFs is likely a direct effect of *csn(lf)* and not a consequence of germline masculinization, as seen with other RNA-binding proteins such as GLD-1 (Jones et al., 1996), as FBF levels were not significantly different in males versus hermaphrodites of the same genetic background (Fig. S4Ai-iv). This function of CSN-5 in FBF protein accumulation, regardless of COP9-dependence, reveals a previously unappreciated role for CSN-5 and also provides the first insight regarding the post-translational regulation of PUF family members FBF-1 and -2. By contrast, we find that CSN-5 does not promote germline stem cell PUF-3 protein accumulation, as seen with FBFs, but might have the opposite effect as we observed a significant increase of PUF-3 accumulation in SPCs of the *csn-5(lf)* mutant germlines (Fig. 4L-N). Recent findings suggest that accumulation of PUF-3 and PUF-11 in the distal mitotic cells is restricted by a TRIM-NHL protein NHL-2 (Brenner et al., 2022) and their degradation at the oocyte to embryo transition is mediated by ubiquitin proteasome system (Spike et al., 2022). It is unclear whether CSN-5 impacts PUF-3 accumulation directly or indirectly and how that might compare to other mechanisms regulating PUF-3 expression. However, the effect on PUF-3 accumulation is distinct from that seen with FBFs indicating CSN-5 specifically regulates FBFs and phenotypes observed in *csn-5(lf)* are not due to a downregulation of multiple PUF proteins.

### CSN-5 promotes FBF protein stability

In this work, we uncovered intrinsic instability of FBF proteins in *csn-5(lf)* mutant background suggesting that *csn-5* promotes FBF accumulation through protecting them from degradation. FBF protein levels in the *csn-5(lf)* genetic background were rescued by disrupting proteasome-mediated degradation without affecting *fbf* transcript levels (Fig. 7). Interestingly, we find that FBF-2, but not FBF-1, levels are partially rescued in the *csn-5(lf)* genetic background with *ned-8(RNAi)* (Fig. 8A-F). This suggests that CSN-5 likely regulates FBF-1 in a COP9-independent manner, whereas regulation of FBF-2 might involve a combination of COP9-dependent and –independent mechanisms. These distinct mechanisms of FBF-1/-2 regulation by CSN-5 are also supported by the observation that FBF-1 levels were most drastically reduced in *csn-5(lf)* whereas FBF-2 levels were equally reduced among all mutants (Fig. 4). As cullins are the only known targets of COP9-mediated deneddylation (Qin et al., 2020), it’s not surprising we have never detected an accumulation of neddylated FBF-2 by Western blot analysis of *csn(lf)* mutant lysates. Although stabilization of FBF-2 requires CSN-5 in the context of COP9, the mechanistic connection between COP9-mediated cullin deneddylation and accumulation of (non-cullin) FBF-2 awaits further study.

### CSN-*5 contribution to germline sex determination*

In this study, we found that *csn-5* contributes to the regulation of germline sex determination through maintenance of FBF protein levels. Loss of function of both *csn-5* and *csn-6* led to a failure to generate oocytes in the majority of mutant germlines (Fig. 6A, B). This loss of oogenic fate was confirmed through a lack of detectable expression of 3xV5::PUF-3 in the proximal pachytene cells of *csn-5(lf)* (Fig. 6D). By contrast, oocytes formed in the majority of *csn-2(lf)* mutant germlines (Fig. 6Aii, B), and this residual oogenesis in *csn-2(lf)* was dependent on FBF-1 as levels of germline masculinization in *csn-2(lf); fbf-1(lf)* increased to those seen in *csn-5(lf)* (Fig. 6B). The function of *csn-5* in germline sex determination appears to be cell-autonomous as the *3xflag::csn-5* transgene driven by the *gld-1* promoter (Ellenbecker et al., 2019) and expected to only be expressed in the germline rescued oocyte formation to 100%, similar to the *3xflag::csn-5* transgene under the control of *csn-5* promoter (Table 1).

Does the mutation of *csn-5* compromise other subunits of the COP9 complex? Previously-published work in human cells, *Drosophila* larvae, and with recombinant proteins suggests that the loss of CSN5 leaves the remainder of COP9 subunits largely intact and assembled, except unable to carry out deneddylation of cullin substrates (Oron et al., 2002; Groisman et al., 2003; Yun et al., 2004; Peth et al., 2007; Sharon et al., 2009). Therefore, we interpret *csn-5(lf)* phenotypes as reflecting the loss of CSN-5 protein and deneddylation, but not affecting the other COP9 subunits. By contrast, loss of several other COP9 subunits results in downregulation of CSN5, as observed for knockdowns of human CSN1 and CSN3 (Peth et al., 2007) as well as for knockdown or mutation of *C. elegans csn-6* (Miller et al., 2009; Fig. S4B). Therefore, care must be taken to distinguish whether any phenotypes of *csn-6(lf)* result from the secondary loss of CSN-5. We approached this by introducing an additional copy of *csn-5* into *csn-6(lf)* genetic background, which was sufficient to rescue oogenesis (Fig. 6C). This suggests that defective oogenesis in *csn-6(lf)* results from the secondary loss of CSN-5. Additionally, this result argued that CSN-5 is sufficient to promote oogenesis in the absence of CSN-5/CSN-6 heterodimer, since *csn-6(lf)* is a null allele where maternal protein stockpile is expected to be depleted (Brockway et al., 2014). Overall, we conclude that CSN-5 is able to promote oogenesis outside of COP9 holoenzyme.

Finally, several lines of evidence suggest that CSN-5 can promote female germ cell fate independent of deneddylating activity. First, *csn-2(lf)* is expected to disrupt COP9 dependent deneddylation, yet it still allows oogenesis in ∼70% of mutant germlines. Second, CSN6 is critically required for COP9-dependent deneddylation (Birol et al., 2014), yet *3xflag::csn-5* rescues oogenesis in *csn-6(lf)* genetic background. Third, *csn-5* can support considerable oogenesis independent of its deneddylating activity as we observe a rescue of fertility to ∼46% in the catalytically inactive, germline-specific, *3xflag::csn-5(D152N); csn-5(lf)* (Table 1). An observation that appears in conflict with this conclusion is the 100% rescue of oogenesis by *ned-8(RNAi)* in *csn-5(lf)* background (Fig. 8H). However, *ned-8(RNAi)* did not affect FBF-1 levels and rescued FBF-2 amounts only in ∼63% germlines (Fig. 8A, B), suggesting that the rescue of oogenesis is independent of FBF levels in at least 37% of germlines. This FBF-independent oogenesis might be explained by a reduction in activity of cullin-based E3 ligase CUL-2/FEM-1/FEM-2/FEM-3 resulting from downregulation of neddylation via *ned-8* knockdown. Neddylation is required for the function of all cullin-based E3 ligases (Duda et al., 2008; Saha and Deshaies, 2008; Boh et al., 2011), and CUL-2/FEM-1/FEM-2/FEM-3 is required for spermatogenesis to degrade TRA-1 (Starostina et al., 2007). The CUL-2/FEM-1/FEM-2/FEM-3 complex functions downstream of FBFs in germline sex determination pathway (Ellis, 2021) and *fem-3* is epistatic to *fbfs* (Zhang et al., 1997). Therefore, it is possible that *ned-8(RNAi)* indirectly circumvents the requirement for *csn-5* in oogenesis and that the contribution of COP9-mediated deneddylation to germline sex determination is through maintaining the balance of neddylated/deneddylated CUL-2/FEM-1/FEM-2/FEM-3.

### CSN-5 role in stem cell maintenance

We find that both CSN-5 and COP9 signalosome are required for germline stem cell maintenance in *C. elegans*. Our findings are consistent with a previous report in *Drosophila* ovarian germline stem cells (Pan et al., 2014), and identify FBFs as a specific target of CSN-5 regulation relevant for stem cell maintenance. In agreement with reduced FBF levels, we observed a significant loss of SPCs in *csn(lf)*, with the most drastic decrease for *csn-5/-6(lf)* (Fig. 5A, B). Although this phenotype is consistent with *fbf(lf)*, it is not exclusive to *fbf(lf)* and could be caused by *csn(lf)* affecting other stem cell regulators. However, we found that *csn-5* functions within the same genetic pathway as *fbfs* when we performed *fbf* knockdown in *csn-5(lf)* mutant and found no further reduction in number of SPCs (Fig. S3A). Additionally, *csn-5(lf)* did not reduce the levels of PUF-3 (Fig. 4L-N), further supporting the notion that it affects stem and progenitor cells though regulation of FBF levels. It is still unclear whether CSN-5’s function in stem cell maintenance depends on the presence of other COP9 subunits as we were unable to rescue SPC numbers in *csn-6(lf)* mutant by providing an extra copy of *3xflag::csn-5*. This result might be due to the insufficient levels of the transgenic protein (Fig. S4B). Alternatively, it remains possible that CSN-5 needs to form a heterodimer with CSN-6 to promote stem cell proliferation. We couldn’t establish whether stabilization of FBFs by inhibiting proteasome-dependent degradation would suppress the SPC reduction in *csn-5(lf)* since *pbs-6(RNAi)* caused reduction of SPC numbers in the control background (data not shown) likely due to its effects on the cell cycle. However, we find a partial rescue of SPC numbers in *csn-5(lf) ned-8(RNAi),* which is consistent with a partial rescue of FBF-2 function, along with a partial rescue of FBF-2 protein levels (Fig. 8B, G). Therefore, we conclude that deneddylation activity of CSN-5 contributes to stem cell proliferation, possibly through regulation of FBF-2 levels. We considered whether *csn-5(lf) ned-8(RNAi)* produced a germline state reminiscent of *fbf-1(lf)* since we observed an increase in FBF-2 protein without a rescue of FBF-1. *fbf-1(lf)* mutation results in a smaller progenitor zone compared to the wild type germlines (Lamont et al., 2004). However, the number of SPCs in *csn-5(lf) ned-8(RNAi)* (∼0.62x of the wild type on average, Fig. 8G) is smaller than that of *fbf-1(lf)* mutant (∼0.85x of the wild type; Wang et al., 2020), suggesting that the partial rescue of FBF-2 levels is not sufficient to recapitulate its full functionality.

A number of previously characterized mutants of splicing factors exhibit a combination of defective female fate specification as well as underproliferated germlines, reminiscent of those observed in *csn-5* and *csn-6* mutants (Puoti and Kimble, 1999, 2000; Belfiore et al., 2004; Konishi et al., 2008; Mantina et al., 2009; Zanetti et al., 2011; Wang et al., 2012). This raised the question of whether underproliferation observed in *csn-5* and - *6(lf)* was linked to the failure of sperm/oocyte switch. However, we could not rescue number of SPCs (data not shown) with *3xflag::csn-5; csn-6(ok1604)* despite being able to rescue oogenesis (Fig. 6C), suggesting that the reduction in SPCs in *csn-6(lf)* does not result from a defective switch to oogenesis. Additionally, male germlines maintain similar SPC population to the hermaphrodite germlines, suggesting no intrinsic sex specific differences in stem cell numbers (Morgan et al., 2010).

The SPCs of *csn-5/-6(lf)* germlines continue to proliferate and enter meiosis (Fig. 5C-G) indicating that the smaller germlines of these genetic backgrounds (Fig. 5A, 6A) are not the result of arrested cell divisions. However, we also observed that *csn-5/-6(lf)* germline SPCs incorporated EdU slower than wild type, requiring longer EdU exposures to label nuclei (data not shown), suggesting a slower cell cycle. Notably, homologs of COP9 complex subunits have previously been found regulating proteins involved in cell cycle progression (Tomoda et al., 1999; Yang et al., 2002; Doronkin et al., 2003; Yoshida et al., 2013), which might provide some explanation behind our observations in the *csn(lf)* mutants. At this time, it remains unknown if the balance between proliferation and differentiation is disrupted in these *csn(lf)* mutants, and further research would be required to understand their cell cycle dynamics. In summary, we conclude that *csn-5* promotes stem cell maintenance through regulation of FBF protein accumulation and speculate this might be a conserved mechanism by which *csn-5* impacts stem cell maintenance in other species.

### Potential mechanisms behind CSN-5 stabilization of FBFs

CSN-5/CSN5 is reported to interact with a number of proteins beyond components of the signalosome (for examples see Tomoda et al., 1999; Bae et al., 2002; Smith et al., 2002; Yoshida et al., 2013) and to promote accumulation of some of its interacting partners (Bemis et al., 2004; Orsborn et al., 2007). Our work defines a novel class of CSN-5 interactors, the PUF-family FBF proteins that are stabilized by CSN-5 and reveals a new mechanism of CSN-5 contribution to stem cell maintenance. Based on our detection of binding between the human homologs, we speculate that CSN5 might similarly stabilize PUM1. The mechanism of CSN-5-dependent stabilization of its partner proteins remains unclear, especially considering there might be several mechanisms at play as revealed by this work. One possible mechanism suggested by previous reports in both fission yeast and mammalian cells is recruitment of a **<u>d</u>**e**<u>ub</u>**iquitinating enzyme (DUB) Ubp12p/USP15 to the COP9 signalosome substrates (Zhou et al., 2003; Hetfeld et al., 2005). Importantly, in several cases DUB recruitment was shown to depend on CSN5 (Groisman et al., 2003; Wee et al., 2005; Liu et al., 2009). Notably, if FBFs are substrates of CSN-5-associated DUBs, a transient *in vivo* association between FBF-1 and CSN-5 would not be surprising. Similarly, recruitment of DUBs would be consistent with the ability of catalytically-dead CSN-5(D152N) to rescue FBF function. Alternatively, CSN5 was reported to directly interact with HIF1α and to stabilize HIF1α in a COP9-independent manner (Bemis et al., 2004). Stabilization of HIF1α was mediated by preventing proline hydroxylation of HIF1α and by interfering with the association of hydroxylated HIF1α with pVHL ubiquitin ligase (Bemis et al., 2004). As a third option, CSN5 metalloprotease was proposed to directly deubiquitinate its binding partner (Lim et al., 2016; Liu et al., 2020). Overall, CSN-5 might be able to protect FBFs from ubiquitination and thus promote their stabilization in *C. elegans* germline stem cells through either of these mechanisms.

CSN-5 contribution to the regulation of germline stem cell proliferation through stabilization of FBF proteins has major implications for understanding regulatory mechanisms relevant for stem cell maintenance. Intrinsic instability of PUF-family proteins counteracted by CSN-5 might reflect a general mechanism regulating the abundance of essential stem cell factors and their effects on stem cell population dynamics. Additionally, the function of CSN-5 in FBF stabilization may be relevant for understanding cancer as CSN5 is upregulated in many human cancers (Lee et al., 2011; Liu et al., 2020). Similarly, a number of cancers overexpress PUM1 and/or PUM2 (Guan et al., 2018; Gor et al., 2021; Shi et al., 2021; Smialek et al., 2021). This PUM protein overexpression in cancer might be borne out through its stabilization via CSN5 and therefore reflect an unappreciated mechanism promoting cancer cell proliferation.

## Materials and Methods

### Nematode strains and culture

All *C. elegans* strains (Table S2) used in this study were cultured on New Nematode Growth Medium (NNGM) plates seeded with *E. coli* OP50 as per standard protocols (Brenner, 1974). c*sn(lf)* mutants used in this study were obtained from the *Caenorhabditis* Genetics Center and outcrossed >8 times to wild type before analysis. The strains were maintained at 20°C, unless specified otherwise.

### csn-2(ok1288) allele

In order to determine the molecular nature of *csn-2(ok1288)* allele, we conducted PCR analysis of genomic DNA as well as cDNA. Total RNA isolated from *csn-2(ok1288)* hermaphrodites was reverse-transcribed using SuperScript IV reverse transcriptase (Invitrogen) and amplified with the primers AGACCCAGGAAAAGTTCGGT and GAGACCATCATCCAAAATTGCGT. Our analysis revealed that the 1680-nt genomic deletion removes most of intron 3, exon 4, and most of intron 4. The *csn-2(ok1288)* transcript is spliced from the end of exon 3 to the start of exon 5 resulting in a frameshift at amino acid 238 and a premature stop codon (removing 50% of ORF; Fig. S6). The full-length CSN-2 contains a PCI domain consisting of helical repeats 1-8 and a winged helix (WH) subdomain, which is followed by a C-terminal alpha helix. CSN-2 is incorporated into the COP9 signalosome through association of the WH subdomain with the WH subdomains of other PCI proteins and through integration of the C-terminal helix into a bundle containing helices of all COP9 subunits (Lingaraju et al., 2014). *ok1288* deletion removes three C-terminal PCI repeats, the WH domain, and the C terminal helix suggesting that the remaining fragment would not incorporate into COP9 signalosome.

### Transgenic animals

All transgene constructs were generated by Gateway cloning (Thermo Fisher Scientific). The two *3xflag::csn-5* transgene constructs contain either the *csn-5* promoter (1110 bp upstream of the CDS) or *gld-1* promoter (1165 bp upstream of the CDS; Ellenbecker et al., 2019) and both contain *csn-5* genomic coding and 3’UTR (733 bp downstream of the CDS) sequences in pCFJ150 (Frokjaer-Jensen et al., 2008). The *3xflag::csn-5(D152N)* transgene construct contains the *gld-1* promoter, *csn-5(D152N)* mutant genomic coding sequence, and *csn-5* 3’UTR sequence in pCFJ150. Mutagenesis of *csn-5(D152N)* was performed using a Q5 site-directed mutagenesis kit per manufacturer instructions (New England BioLabs). For transgenic lines used in PLA, *3xflag::csn-5* was crossed with pre-existing *patcGFP::fbf* or *patcGFP* lines (Wang et al., 2020). For all *csn-5* transgenes, a single-copy insertion of the transgene was generated by homologous recombination into the universal *Mos1* insertion site on LG II after Cas9 induced double-stranded break (Dickinson et al., 2013; Wang et al., 2016). Transgene insertion was confirmed by PCR.

### Yeast Two-Hybrid library screening and directed assays

For the yeast two-hybrid screen, *fbf-2* cDNA was cloned into the pGBTK vector (Clontech) and transformed into PJ68-4a yeast strain (*MAT**a** trp1-901 leu2-3,112 ura3-52 his3-200 gal4Δ gal80Δ LYS2::GAL1-HIS3 GAL2-ADE2 met2::GAL7-lacZ*; James et al., 1996). The yeast two-hybrid library was generated from the poly(A)+ mRNA isolated from mixed-stage N2 nematodes using the Matchmaker kit (Clontech) as per the manufacturer’s protocol and transformed into Y1HGold strain (*MAT**α**, ura3-52, his3-200, ade2-101, trp1-901, leu2-3, 112, gal4Δ, gal80Δ, met–, MEL1*). The bait and prey library strains were mated and cultured on selective plates SD/-Trp, Leu, His, Ade with 1 mM 3-AT; the total number of screened diploids was estimated at 2.32x10^8^. Out of 271 positive colonies, 99 (37%) were identified as *csn-5* fragments by PCR and sequencing of prey plasmid inserts. For directed interaction assays, FBF-1 or FBF-2 in a bait vector pGBKT7 and CSN-5 in a prey vector pGADT7 were cotransformed in PJ69-4a. Empty prey vector was used as the control. Expression of c-myc tagged FBF-1 or FBF-2 and HA tagged CSN-5 proteins in yeast colonies were confirmed by Western blot, see Table S3 for antibody information. Serially diluted yeast cultures (at OD600 0.2) expressing *fbf* bait and *csn-5* prey (or empty prey) were spotted on control or the interaction selection plate SD/-Trp, -Leu, -His, -Ade (with 1mM 3-AT to inhibit leaky expression of Histidine reporter) and incubated at 30°C for 4 days.

### GST pulldown assay

Expression constructs containing His_6_::FBF-1 and His_6_::FBF-2 have been described before (Wang et al., 2016). Truncated FBF expression constructs were generated by PCR from full-length FBF-1 or FBF-2 and inserted into pETDuet-1 plasmid (EMD Millipore). Full-length *C. elegans* CSN-5 (amino acids 1-368) and CSN-6^MPN^ (amino acids 1-197) were amplified from N2 Bristol cDNA and cloned into pGEX-KG plasmid. Mutant CSN-5(D152N) was generated from full-length CSN-5 using a Q5 site-directed mutagenesis kit per manufacturer’s instructions (New England BioLabs). Truncated CSN-5 constructs were made by PCR from full-length CSN-5 and inserted into pGEX-KG plasmids. Human CSN5^MPN^ (amino acids 1-257) was amplified from HEK293 cDNA and cloned into pGEX-KG plasmid. Human PUM1^RBD^ (isoform 1, amino acids 828-1176) and PUM2^RBD^ (isoform 3, amino acids 706-1055) were amplified from HEK293 cDNA and cloned into pETDuet-1 plasmid. All constructs were sequenced and transformed into *Escherichia coli* strain BL21(DE3) for expression. Expression of His-tagged FBF protein constructs was induced with 0.1 mM IPTG at 15°C for 20-24 hours. Expression of GST alone or GST-tagged CSN protein constructs was induced with 0.1 mM isopropyl β-D-1-thiogalactopyranoside (IPTG) at 37°C for 4.5 hours. Pellets of induced bacteria were collected and resuspended in lysis buffer (20 mM Tris pH 7.5, 250 mM NaCl, 10% glycerol, 1 mM MgCl_2_, 0.1% Triton X-100) containing 10 mM beta mercaptoethanol (BME), 1 mM phenylmethylsulfonyl fluoride (PMSF), 1x Roche protease inhibitor cocktail, 0.5 mg/ml lysozyme, and 6 µg/ml DNase I, rotated at 4°C for 1 hour to lyse, and cleared by centrifugation at 1110 *g* for 20 min. Cell lysates of GST alone or GST-tagged CSN constructs were bound to glutathione beads in 20 mM Tris pH 7.5, 250 mM NaCl, 10% glycerol, 0.1% Triton X-100, 10 mM BME, 1 mM PMSF, and 1x Roche protease inhibitor cocktail for 1 hour at 15°C. Binding reactions with His_6_ tagged preys were incubated at 15°C for 3 hours and washed four times with 10 mM Tris pH 7.5, 150 mM NaCl, 0.1% NP-40, and 0.5 mg/ml BSA. For elution, beads were heated to 95°C for 5 minutes in sodium dodecyl sulfate (SDS) sample buffer and 10 mM dithiothreitol (DTT). To test if the FBF-2/CSN-5/-6 binding interaction is RNA-dependent, 50 µg/ml RNase A was added to His_6_::FBF-2 lysate before incubation with GST alone or GST::CSN-5/-6.

### Western blot analysis

Protein samples from pulldown assays were separated using Mini-PROTEAN TGX 4-20% precast gels (Bio-Rad,) and visualized using either Coomassie, stain-free chemistry, or by Western blotting. For anti-His probing, all eluates were treated the same. For anti-GST probing, GST negative control eluate was diluted 8-fold further than GST::CSN constructs to avoid overexposure. See Table S3 for antibody information. Expression, and attachment to GSH beads, of all GST alone and GST-tagged constructs was confirmed for each pulldown experiment.

To analyze FBF protein levels in *csn(lf)* mutants, 50-100 young adult worms of each genotype were picked from synchronous cultures, and lysed in SDS-PAGE sample buffer containing 2.8% SDS and 75mM DTT by boiling for 15 minutes prior to SDS PAGE gel electrophoresis. Protein samples were separated on a 7.5% or 4-20% gel (Bio-Rad) and transferred to a 0.2 µm PVDF membrane (EMD Millipore). After transfer, membranes were blocked in 50 mM Tris pH 7.5, 180 mM NaCl, 0.05% Tween 20 with 5% non-fat dry milk powder, and probed with antibodies (see Table S3) diluted in blocking solution. Membranes were developed using Luminata Crescendo Western HRP substrate (EMD Millipore) and visualized using Bio-Rad ChemiDoc™ MP Imaging System. Band intensities were quantified using Image Lab software version 5.1.

### Proximity ligation assay (PLA)

Duolink® PLA was performed on dissected *C. elegans* gonads following a modified PLA protocol as described (Day et al., 2020; Wang et al., 2020). Fixation was as described below in “Immunofluorescence” section. Blocking steps included incubation in 1xPBS/0.1% Triton-X-100/0.1% BSA (PBS-T/BSA) for 2×15 minutes at room temperature, in 10% normal goat serum for 1 hour at room temperature in a light-sealed humid chamber, and in Duolink blocking buffer (Sigma-Aldrich) for 1 hour at 37°C in a light-sealed humid chamber. Primary antibodies (see Table S3 for antibody information) were diluted in PBS-T/BSA and incubated overnight at 4°C. Samples were then incubated with Duolink PLA probes (see Table S3) for 1 hour at 37°C in a light-sealed chamber. Next, slides were incubated in a light-sealed humid chamber for 30 minutes at 37°C for ligation, and then for 100 minutes at 37°C for amplification before finally being mounted with Duolink Mounting medium with DAPI (Sigma-Aldrich). Images were acquired using Zeiss 880 confocal microscope, and the ImageJ “Analyze Particles” plug-in was used to quantify PLA foci in germline images.

### RNA extraction and RT-qPCR

*C. elegans* were synchronized by bleaching and hatched L1 larvae were plated on NNGM plates with OP50 bacteria, grown at 20°C, and collected after 72 hours. 200-400 worms were collected per biological replicate and stored in Trizol (Invitrogen) at -80°C. Total RNA was extracted using either Monarch Total RNA Miniprep Kit (New England Biolabs) or Direct-zol™ RNA MiniPrep kit (Zymo Research) as per manufacturers’ protocols. RNA concentration was measured using Qubit (Thermo Fisher). cDNA was synthesized using the SuperScript™ IV reverse transcriptase (Invitrogen) using 300 ng RNA template per each 20 µL cDNA synthesis reaction. Quantitative PCR reactions were performed in technical triplicates per each input cDNA using iQ SYBR® Green Supermix (Bio-Rad). Primers for all targets were as previously described: *act-1* and *fbf-2* (Chauve et al., 2020), *unc-54* (Wang et al., 2020), and *fbf-1* (Voronina and Seydoux, 2010), all primers are listed in Table S4. Abundance of each mRNA in *csn(lf)* mutant relative to the wild type was calculated using comparative ΔΔCt method (Plaffl, 2001) with actin *act-1* as a reference gene. After mRNA abundance of each tested gene was normalized to *act-1*, the fold change values from replicates were averaged.

### Immunofluorescence

Adult *C. elegans* hermaphrodites were washed in M9 and germlines were dissected on poly-L-lysine treated slides, flash frozen, and fixed for 1 minute in 100% methanol (- 20°C) followed by 5 minutes in 2% paraformaldehyde/100 mM K_2_HPO_4_ (pH 6) at room temperature. The samples were then blocked in PBS-T/BSA for 30 minutes at room temperature. Next, samples were incubated with primary antibody (see Table S3 for antibody information) diluted in PBS-T/BSA overnight at 4°C. Samples were then washed with PBS-T/BSA three times for 10 minutes per wash, before incubating with secondary antibody diluted in PBS-T/BSA for 2 hours at room temperature in a dark humid chamber. Samples were washed again three times for 10 minutes per wash before adding 10 µL Vectashield® with DAPI (Vector Laboratories) and cover-slipping. Images were acquired with a Leica DFC300G camera attached to a Leica DM5500B microscope with a 40x LP FLUOTAR NA1.3 objective using LAS X software (Leica) and stitched together using Adobe Photoshop CS3.

### Fluorescence quantitation

To analyze PUF-3 levels in *csn-5(lf)* and FBF levels in *csn-5(lf)* mutant treated with control or *ned-8(RNAi)*, adult *C. elegans* hermaphrodites were washed in M9, germlines were dissected, and slides were processed as described above under the “Immunofluorescence” section. Z-stack images spanning the thickness of germline were taken under identical conditions across all samples in each experiment. PUF-3 and FBF protein levels within the SPC zone were analyzed using Fiji/ImageJ as previously described with some modifications (Brenner and Schedl, 2016; Haupt et al., 2019). On the maximum intensity projection generated from each z-stack, the segmented line (width set to 45) tool was used to draw a freehand line from the distal tip through the length of the SPC zone (as indicated by co-staining with REC-8 antibody, Hansen et al., 2004 or DNA morphology with DAPI) that bisected the gonad. Next, pixel intensity data for both V5 and FBF-1 channels were obtained using “Plot Profile” to generate raw intensity curves. To adjust for non-specific background staining, we subtracted the average intensity of the respective negative controls (N2 for V5::PUF-3 analysis and *fbf-1(ok91)* for FBF-1 and V5::FBF-2 analysis) from expression curves of each respective genotype. The maximum background-subtracted values for each wild type and mutant germline were determined and used in further analysis. The maximum values of wild type germlines were averaged, and the average value was used for normalization of all values to set the wild type levels to 1. To better visualize 3xV5::PUF-3 expression in the SPC zone for Fig. 4L, M, images were acquired using a Zeiss 880 confocal microscope under identical conditions and V5 intensity was adjusted equally in Adobe Photoshop CS3 for *3xv5::puf-3* and *csn-5(lf) 3xv5::puf-3* genetic backgrounds.

### Germline SPC counts

*C. elegans* were synchronized by bleaching and hatched L1 larvae were plated on NNGM plates with OP50 bacteria, grown at 20°C, and harvested 24 hours post-L4. Gonads were dissected and stained for progenitor cell marker REC-8 (Hansen et al., 2004), and the number of stem and progenitor cells was measured by counting the total number of cells positive for REC-8 staining using Cell Counter plug-in in Fiji (Schindelin et al., 2012) or Imaris 9.9 software.

### EdU labeling

EdU labeling was performed as previously described with some modifications (Wang et al., 2020). EdU bacterial plates were prepared by diluting an overnight culture of thymine deficient MG1693 *E. coli* (The Coli Genetic Stock Center; Yale University) 1/25 in 1% glucose, 1 mM MgSO_4_, 5 µg/mL thymine, 6 µM thymidine, and 20 µM EdU in M9. This culture was grown at 37°C for 24 hours, pelleted by centrifugation, resuspended in 10 mL M9 and plated on NNGM plates. Worm strains were synchronized by bleaching and hatched L1 larvae were plated on NNGM plates with OP50 bacteria, grown at 20°C for 72 hours to reach young adult stage when they were exposed to EdU-labeled bacteria for either 4 or 14 hours (Fig. 5C, D and E-G, respectively). After feeding for the specified time, worms were dissected for immunostaining as described in “Immunofluorescence” section above. After incubation with secondary antibody (see Table S3 for antibody information), slides were washed four times for 15 minutes per wash in PBS-T/BSA. Next, the Click-iT reaction was performed according to manufacturer instructions with the exception that two 30-minute Click-iT reactions were performed to increase the signal of the Alexa 488 dye. After incubation with the second Click-iT reaction, slides were washed four times for 15 minutes per wash in PBS-T/BSA before adding 10 µL Vectashield® with DAPI (Vector Laboratories) and cover-slipping.

### Phenotypic analysis

For germline sex analysis (Fig. 6), *C. elegans* were synchronized by bleaching and hatched L1 larvae were plated on NNGM plates with OP50 bacteria, grown at 20°C, and harvested after 24 hours post-L4. Gonads were dissected, stained for DNA with DAPI, and scored for abnormal oocytes (defined as <3 oocytes per germline, uncondensed chromosomes, endomitotic oocytes, or increased number of DAPI spots indicating achiasmatic chromosomes), sperm only, and failure of gamete formation.

For scoring fertility in *csn-5(ok1064)* mutant strains (Table 1), *C. elegans* were synchronized by bleaching and hatched L1 larvae were plated on NNGM plates with OP50 bacteria and grown at 20°C for 96 hours before scoring fertility using a dissecting microscope. “Fertile” was classified by presence of embryos in the uteri and lack of embryos was classified as sterile.

### RNAi treatment

For RNAi, two knockdown methods were utilized. Feeding RNAi was utilized for *pbs-6* and *fbf-2* RNAi constructs in pL4440 (Kamath and Ahringer, 2003). Empty vector pL4440 was used as a control in all RNAi experiments. All RNAi constructs were verified by sequencing and transformed into HT115 (DE3) *Escherichia coli*. Three colonies of freshly transformed RNAi plasmids were combined for growth in LB/75 µg/mL carbenicillin media for 4 hours, and induced with 5 mM IPTG for 1-2 hours at 37°C. RNAi plates (NNGM containing 100 µg/mL carbenicillin and 0.4 mM IPTG) were seeded with the pelleted cells. Synchronously hatched L1 larvae were plated directly on RNAi plates, grown at 24°C, and collected for analysis after reaching adulthood.

Soaking RNAi for *ned-8* was performed as described by Yoon et al., 2012 with some modifications. Templates for *in vitro* transcription of *ned-8* flanked by T7 promoter sequences were amplified via PCR from RNAi constructs in pL4440 (Source BioScience RNAi library; Kamath and Ahringer, 2003). Templates were then transcribed into dsRNA *in vitro* using a T7 MEGAscript® Kit (Invitrogen) per manufacturer’s instructions at the 100 µL scale. dsRNA was then recovered using an RNA Cleanup Kit (New England Biolabs) per manufacturer’s instructions. Synchronously hatched L1 larvae (200-400) were then suspended in soaking buffer containing 2.5X M9, 30 mM spermidine, and 0.5% gelatin with 1 µg/µL dsRNA for 48 hours at 20°C. Larvae were then plated on OP50 plates for another 72 hours, and were collected for analysis after reaching adulthood.

## Acknowledgements

We thank members of the Voronina laboratory for helpful discussions. All research was performed at the University of Montana. Several nematode strains were provided by the *Caenorhabditis* Genetics Center, funded by the National Institutes of Health (NIH) (grant P40OD010440). *csn* knockout strains were generated by the *C. elegans* Reverse Genetics Core Facility at the University of British Columbia and Oklahoma Medical Research Foundation, both part of the international *C. elegans* Gene Knockout Consortium (Barstead et al., 2012). Confocal microscopy was performed in the University of Montana BioSpectroscopy Core Research Laboratory operated with support from NIH awards P20GM103546 and S10OD021806. We are grateful to Gabriella Weiss for help with strain generation, Nicholas Day for assistance with Western blot, and James Bosco, Ella Baumgarten, and Polash Biswas for help with cloning. We thank Beverly Piggott and Isabella Maag for sharing, and instruction, of Imaris cell counting software.

## Author Contributions

E.O. contributed to all Figures, participated in study design, manuscript drafting and editing. M.E. provided strains, constructs, and experimental data related to Fig. 2, 4-6, S1, S2, and S6. X.W. provided constructs and contributed to Fig. 1, 2, 4, and Table S1. M.T., K.J., and D.C. contributed strains and experimental data related to Fig. 5-7, S4, S5, and Table 1. E.V. contributed to generation of strains and plasmids, provided general study design, coordination, manuscript drafting and editing. All authors made manuscript revisions and approved the manuscript.

## Data availability

This study did not generate large datasets or codes, but raw data/images are available upon request.

---

**Fig. S1. RNA-independent FBF-2 interaction with CSN-5 and CSN-6.**

(A) GST pulldown assay of His_6_::FBF-2 with GST::CSN-5^MPN^ or (B) GST::CSN-6^MPN^ with or without RNase A treatment. FBF-2 protein is detected by Western blot with anti-His. Total protein seen by Coomassie.

**Fig. S2. Relative *fbf-2* transcript abundance is reduced in *csn(lf)* mutants.**

Steady-state levels of transcripts indicated on the X-axis in *3xv5::fbf-2(q932)*, referred to here as wild type, and *3xv5::fbf-2(q932); csn(lf)* mutants are quantified by RT-qPCR and normalized to reference gene *act-1*. *unc-54* is a control. Differences in relative mRNA abundance of *fbf-1/-2* was evaluated by one-way ANOVA with Dunnett’s post-test. Grey asterisks denote statistical significance compared to wild type (****, *p*<0.0001). Data are representative of five biological replicates of wild type, four replicates of both *csn-2(lf)* and *csn-6(lf)*, and three biological replicates of *csn-5(lf)*. Data are plotted as mean values with error bars representing standard deviation.

**Fig. S3. *csn-5(lf)* functions in the same pathway as *fbfs* in SPC maintenance.**

Quantification of SPCs (by anti-REC-8 staining) after treatment with a control RNAi (circles) and *fbf(RNAi)* (triangles) in *3xv5::fbf-2(q932); csn-5(lf)/nT1*, referred to here as *csn-5(ok1064)/nT1* (grey), and *3xv5::fbf-2(q932); csn-5(lf)* referred to as *csn-5(ok1064)* (red). Differences in number of SPCs were evaluated by one-way ANOVA with Sidak’s post-test, ns denotes not significant. Number of germlines scored (*N*) are indicated at the bottom of the graph. Data reflects two biological replicates.

**Fig. S4. FBF levels are similar in hermaphrodites and males; CSN-5 levels are reduced in *csn-6(lf)*, and catalytically dead CSN-5 maintains binding to FBF-2 *in vitro*.**

(Ai, iii) Western blot analysis comparing worm lysate of hermaphrodites to males in *rrf-1(pk1417); 3xv5::fbf-2(q932); him-8(tm611)* mutants. Endogenous FBF-1 and epitope tagged endogenous 3xV5::FBF-2 are detected by anti-FBF-1 and anti-V5. Tubulin is used as a loading control. (Aii, iv) Total protein in males was normalized to hermaphrodites. Differences in protein level were evaluated by Student’s t-test, ns denotes not significant. The data reflect four biological replicates for FBF-2 and three biological replicates for FBF-1. Mean group values are shown as lines and error bars denote standard deviation. (Bi) 3xFLAG::CSN-5 protein levels in *3xflag::csn-5* and *3xflag::csn-5; csn-6(ok1604)* animals detected by Western blot with anti-FLAG. Tubulin is used as a loading control. (Bii) Relative level of 3xFLAG::CSN-5 protein in *csn-6(lf)* was normalized to the wild type. Difference in protein level was evaluated by Student’s t-test. Asterisks denote statistical significance (***, *p*<0.005). The data reflect six biological replicates, is represented as average values, and error bars denote standard deviation. (C) GST pulldown of His_6_::FBF-2 with GST::CSN-5(D152N). Protein constructs are detected by Western blot with anti-His and anti-GST, the positions of GST::CSN-5 constructs are marked by a black dot.

**Fig. S5. Oogenesis is restored in *rrf-1(lf); csn-5(lf)* mutant backgrounds.**

Gametogenesis in *pbs-6(RNAi)-*treated *rrf-1(pk1417); fbf-2(q932)* and *rrf-1(pk1417); fbf-2(q932); csn-5(ok1064)* mutants. Number of germlines scored (*N*) are indicated at the top of each column. Data are plotted in aggregate and reflect two biological replicates.

**Fig. S6. *csn-2(ok1288)* allele used in this study.**

(A) Schematic diagram of *csn-2(ok1288)* locus with exons pictured as black rectangles, introns as black lines, and the UTRs as grey rectangles. The 1680 nucleotide *ok1288* deletion (red) results in the removal of most of intron 3, exon 4, and most of intron 4. The spliced transcript results in a frameshift at amino acid 238 and a premature stop codon. (B) The schematic of full-length CSN-2 protein compared to CSN-2(ok1288) shows the *ok1288* deletion removing three C-terminal PCI repeats (squares), the winged-helix (WH) subdomain, and the C-terminal helix (oval). Domains of CSN-2 are delineated by homology with human CSN2 from a previous report (Lingaraju et al., 2014).-

## References

1. Ariz, M., Mainpal, R., and Subramaniam, K. (2009). *C. elegans* RNA-binding proteins PUF-8 and MEX-3 function redundantly to promote germline stem cell mitosis. Dev Biol, 326, 295–304.

2. Austin, J., and Kimble, J. (1987). *glp-1* is required in the germ line for regulation of the decision between mitosis and meiosis in *C. elegans*. Cell, 51, 589–599.

3. Bae, M.K., Ahn, M.Y., Jeong, J.W., Bae, M.H., Lee, Y.M., Bae, S.K., Park, J.W., Kim, K.R., and Kim, K.W. (2002). Jab1 interacts directly with HIF-1alpha and regulates its stability. J Biol Chem, 277, 9–12.

4. Barstead, R., Moulder, G., Cobb, B., Frazee, S., Henthorn, D., Holmes, J., Jerebie, D., Landsdale, M., Osborn, J., Pritchett, C. et al. (2012). Large-scale screening for targeted knockouts in the *Caenorhabditis elegans* genome. G3 (Bethesda), 2, 1415–1425.

5. Belfiore, M., Pugnale, P., Saudan, Z., and Puoti, A. (2004). Roles of the *C. elegans* cyclophilin-like protein MOG-1 in MEP-1 binding and germline fates. Development, 131, 2935–2945.

6. Bemis, L., Chan, D.A., Finkielstein, C.V., Qi, L., Sutphin, P.D., Chen, X., Stenmark, K., Giaccia, A.J., and Zundel, W. (2004). Distinct aerobic and hypoxic mechanisms of HIF-alpha regulation by CSN5. Genes Dev, 18, 739–744.

7. Bernstein, D., Hook, B., Hajarnavis, A., Opperman, L., and Wickens, M. (2005). Binding specificity and mRNA targets of a *C. elegans* PUF protein, FBF-1. RNA, 11, 447–458.

8. Berry, L.W., Westlund, B., and Schedl, T. (1997). Germ-line tumor formation caused by activation of *glp-1*, a *Caenorhabditis elegans* member of the *Notch* family of receptors. Development, 124, 925–936.

9. Birol, M., Enchev, R.I., Padilla, A., Stengel, F., Aebersold, R., Betzi, S., Yang, Y., Hoh, F., Peter, M., Dumas, C. et al. (2014). Structural and biochemical characterization of the Cop9 signalosome CSN5/CSN6 heterodimer. PLoS One, 9, e105688.

10. Boh, B.K., Smith, P.G., and Hagen, T. (2011). Neddylation-induced conformational control regulates cullin RING ligase activity *in vivo*. J Mol Biol, 409,136–145.

11. Brenner, J.L., Jyo, E.M., Mohammad, A., Fox, P., Jones, V., Mardis, E., Schedl, T., and Maine, E.M. (2022). TRIM-NHL protein, NHL-2, modulates cell fate choices in the *C. elegans* germ line. Dev Biol, 491, 43–55.

12. Brenner, J.L., and Schedl, T. (2016). Germline stem cell differentiation entails regional control of cell fate regulator GLD-1 in *Caenorhabditis elegans*. Genetics, 202, 1085–1103.

13. Brenner, S. (1974). The genetics of *Caenorhabditis elegans*. Genetics, 77, 71–94.

14. Brockway, H., Balukoff, N., Dean, M., Alleva, B., and Smolikove, S. (2014). The CSN/COP9 signalosome regulates synaptonemal complex assembly during meiotic prophase I of *Caenorhabditis elegans*. PLoS Genet, 10, e1004757.

15. Buck, S.H., Chiu, D., and Saito, R.M. (2009). The cyclin-dependent kinase inhibitors, *cki-1* and *cki-2*, act in overlapping but distinct pathways to control cell cycle quiescence during *C. elegans* development. Cell Cycle, 8, 2613–2620.

16. Chamovitz, D.A. (2009). Revisiting the COP9 signalosome as a transcriptional regulator. EMBO Rep, 10, 352–358.

17. Chamovitz, D.A., Wei, N., Osterlund, M.T., von Arnim, A.G., Staub, J.M., Matsui, M., and Deng, X.W. (1996). The COP9 complex, a novel multisubunit nuclear regulator involved in light control of a plant developmental switch. Cell, 86, 115– 121.

18. Chauve, L., Le Pen, J., Hodge, F., Todtenhaupt, P., Biggins, L., Miska, E.A., Andrews, S., and Casanueva, O. (2020). High-Throughput Quantitative RT-PCR in Single and Bulk C. elegans Samples Using Nanofluidic Technology. J Vis Exp, 159, e61132

19. Chen, J., Mohammad, A., Pazdernik, N., Huang, H., Bowman, B., Tycksen, E., and Schedl, T. (2020). GLP-1 Notch-LAG-1 CSL control of the germline stem cell fate is mediated by transcriptional targets *lst-1* and *sygl-1*. PLoS Genet, 16, e1008650.

20. Claret, F.X., Hibi, M., Dhut, S., Toda, T., and Karin, M. (1996). A new group of conserved coactivators that increase the specificity of AP-1 transcription factors. Nature, 383, 453–457.

21. Cope, G.A., Suh, G.S., Aravind, L., Schwarz, S.E., Zipursky, S.L., Koonin, E.V., and Deshaies, R.J. (2002). Role of predicted metalloprotease motif of Jab1/Csn5 in cleavage of Nedd8 from Cul1. Science, 298, 608–611.

22. Crittenden, S.L., Bernstein, D.S., Bachorik, J.L., Thompson, B.E., Gallegos, M., Petcherski, A.G., Moulder, G., Barstead, R., Wickens, M., and Kimble, J. (2002). A conserved RNA-binding protein controls germline stem cells in *Caenorhabditis elegans*. Nature, 417, 660–663.

23. Day, N.J., Wang, X., and Voronina, E. (2020). *In situ* detection of ribonucleoprotein complex assembly in the *C. elegans* germline using proximity ligation assay. J Vise Exp, 159, e60982.

24. Demarco, R.S., Stack, B.J., Tang, A.M., Voog, J., Sandall, S.L., Southall, T.D., Brand, A.H., and Jones, D.L. (2022). Escargot controls somatic stem cell maintenance through the attenuation of the insulin receptor pathway in *Drosophila*. Cell Rep, 39, 110679.

25. Dessau, M., Halimi, Y., Erez, T., Chomsky-Hecht, O., Chamovitz, D.A., and Hirsch, J.A. (2008). The *Arabidopsis* COP9 signalosome subunit 7 is a model PCI domain protein with subdomains involved in COP9 signalosome assembly. Plant Cell, 20, 2815–2834.

26. Dickinson, D.J., Ward, J.D., Reiner, D.J., and Goldstein, B. (2013). Engineering the *Caenorhabditis elegans* genome using Cas9-triggered homologous recombination. Nat Methods, 10, 1028–1034.

27. Doronkin, S., Djagaeva, I., and Beckendorf, S.K. (2003). The COP9 signalosome promotes degradation of Cyclin E during early *Drosophila* oogenesis. Dev Cell, 4, 699–710.

28. Duda, D.M., Borg, L.A., Scott, D.C., Hunt, H.W., Hammel, M., and Schulman, B.A. (2008). Structural insights into NEDD8 activation of cullin-RING ligases: conformational control of conjugation. Cell, 134, 995–1006.

29. Echalier, A., Pan, Y., Birol, M., Tavernier, N., Pintard, L., Hoh, F., Ebel, C., Galophe, N., Claret, F.X., and Dumas, C. (2013). Insights into the regulation of the human COP9 signalosome catalytic subunit, CSN5/Jab1. Proc Natl Acad Sci U S A, 110, 1273–1278.

30. Eckmann, C.R., Kraemer, B., Wickens, M., and Kimble, J. (2002). GLD-3, a bicaudal-C homolog that inhibits FBF to control germline sex determination in *C. elegans*. Dev Cell, 3, 697–710.

31. Ellenbecker, M., Osterli, E., Wang, X., Day, N.J., Baumgarten, E., Hickey, B., and Voronina, E. (2019). Dynein light chain DLC-1 facilitates the function of the germline cell fate regulator GLD-1 in *Caenorhabditis elegans*. Genetics, 211, 665–681.

32. Ellis, R.E. (2022). Sex determination in nematode germ cells. Sex Dev, 1–18.

33. Enchev, R.I., Scott, D.C., da Fonseca, P.C., Schreiber, A., Monda, J.K., Schulman, B.A., Peter, M., and Morris, E.P. (2012). Structural basis for a reciprocal regulation between SCF and CSN. Cell Rep, 2, 616–627.

34. Fox, P.M., Vought, V.E., Hanazawa, M., Lee, M.H., Maine, E.M., and Schedl, T. (2011). Cyclin E and CDK-2 regulate proliferative cell fate and cell cycle progression in the *C. elegans* germline. Development, 138, 2223–2234.

35. Friend, K., Campbell, Z.T., Cooke, A., Kroll-Conner, P., Wickens, M.P., and Kimble, J. (2012). A conserved PUF-Ago-eIF1A complex attenuates translational elongation. Nat Struct Mol Biol, 19, 176–183.

36. Frøkjaer-Jensen, C., Davis, M.W., Hopkins, C.E., Newman, B.J., Thummel, J.M., Olesen, S.P., Grunnet, M., and Jorgensen, E.M. (2008). Single-copy insertion of transgenes in *Caenorhabditis elegans*. Nat Genet, 40, 1375–1383.

37. Gor, R., Sampath, S.S., Lazer, L.M., and Ramalingam, S. (2021). RNA binding protein PUM1 promotes colon cancer cell proliferation and migration. Int J Biol Macromol, 174, 549–561.

38. Groisman, R., Polanowska, J., Kuraoka, I., Sawada, J., Saijo, M., Drapkin, R., Kisselev, A.F., Tanaka, K., and Nakatani, Y. (2003). The ubiquitin ligase activity in the DDB2 and CSA complexes is differentially regulated by the COP9 signalosome in response to DNA damage. Cell, 113, 357–367.

39. Guan, X., Chen, S., Liu, Y., Wang, L.L., Zhao, Y., and Zong, Z.H. (2018). PUM1 promotes ovarian cancer proliferation, migration and invasion. Biochem Biophys Res Commun, 497, 313–318.

40. Hallstrom, T.C., and Nevins, J.R. (2006). Jab1 is a specificity factor for E2F1 induced apoptosis. Genes Dev, 20, 613–623.

41. Hansen, D., Hubbard, E.J., and Schedl, T. (2004). Multi-pathway control of the proliferation versus meiotic development decision in the *Caenorhabditis elegans* germline. Dev Biol, 268, 342–357.

42. Hansen, D., and Schedl, T. (2013). Stem cell proliferation versus meiotic fate decision in *Caenorhabditis elegans*. Adv Exp Med Biol, 757, 71–99.

43. Haupt, K.A., Enright, A.L., Ferdous, A.S., Kershner, A.M., Shin, H., Wickens, M., and Kimble, J. (2019). The molecular basis of LST-1 self-renewal activity and its control of stem cell pool size. Development, 146, dev181644.

44. Haupt, K.A., Law, K.T., Enright, A.L., Kanzler, C.R., Shin, H., Wickens, M., and Kimble, J. (2020). A PUF hub drives self-renewal in *Caenorhabditis elegans* germline stem cells. Genetics, 214, 147–161.

45. Hetfeld, B.K., Helfrich, A., Kapelari, B., Scheel, H., Hofmann, K., Guterman, A., Glickman, M., Schade, R., Kloetzel, P.M., and Dubiel, W. (2005). The zinc finger of the CSN-associated deubiquitinating enzyme USP15 is essential to rescue the E3 ligase Rbx1. Curr Biol, 15, 1217–1221.

46. Hofmann, K., and Bucher, P. (1998). The PCI domain: a common theme in three multiprotein complexes. Trends Biochem Sci, 23, 204–205.

47. Hubbard, E.J. (2007). *Caenorhabditis elegans* germ line: a model for stem cell biology. Dev Dyn, 236, 3343–3357.

48. James, P., Halladay, J., and Craig, E.A. (1996). Genomic libraries and a host strain designed for highly efficient two-hybrid selection in yeast. Genetics, 144, 1425–1436.

49. Jones, A.R., Francis, R., and Schedl, T. (1996). GLD-1, a cytoplasmic protein essential for oocyte differentiation, shows stage- and sex-specific expression during *Caenorhabditis elegans* germline development. Dev Biol, 180, 165–183.

50. Kalchhauser, I., Farley, B.M., Pauli, S., Ryder, S.P., and Ciosk, R. (2011). FBF represses the Cip/Kip cell-cycle inhibitor CKI-2 to promote self-renewal of germline stem cells in *C. elegans*. EMBO J, 30, 3823–3829.

51. Kamath, R.S., and Ahringer, J. (2003). Genome-wide RNAi screening in *Caenorhabditis elegans*. Methods, 30, 313–321.

52. Kimble, J., and Crittenden, S.L. (2007). Controls of germline stem cells, entry into meiosis, and the sperm/oocyte decision in *Caenorhabditis elegans*. Annu Rev Cell Dev Biol, 23, 405–433.

53. Kimble, J.E., and White, J.G. (1981). On the control of germ cell development in *Caenorhabditis elegans*. Dev Biol, 81, 208–219.

54. Kocsisova, Z., Mohammad, A., Kornfeld, K., and Schedl, T. (2018). Cell cycle analysis in the *C. elegans* germline with thymidine analog EdU. J Vis Exp, 140, e58339.

55. Konishi, T., Uodome, N., and Sugimoto, A. (2008). The *Caenorhabditis elegans* DDX-23, a homolog of yeast splicing factor PRP28, is required for the sperm oocyte switch and differentiation of various cell types. Dev Dyn, 237, 2267–2277.

56. Kraemer, B., Crittenden, S., Gallegos, M., Moulder, G., Barstead, R., Kimble, J., and Wickens, M. (1999). NANOS-3 and FBF proteins physically interact to control the sperm-oocyte switch in *Caenorhabditis elegans*. Curr Biol, 9, 1009–1018.

57. Kumsta, C., and Hansen, M. (2012). *C. elegans rrf-1* mutations maintain RNAi efficiency in the soma in addition to the germline. PLoS One, 7, e35428.

58. Lamont, L.B., Crittenden, S.L., Bernstein, D., Wickens, M., and Kimble, J. (2004). FBF-1 and FBF-2 regulate the size of the mitotic region in *C. elegans* germline. Dev Cell, 7, 697–707.

59. Lee, C., Sorensen, E.B., Lynch, T.R., and Kimble, J. (2016). *C. elegans* GLP 1/Notch activates transcription in a probability gradient across the germline stem cell pool. eLife, 5, e18370.

60. Lee, M.H., Zhao, R., Phan, L., and Yeung, S.C. (2011). Roles of COP9 signalosome in cancer. Cell Cycle, 10, 3057–3066.

61. Lim, S.O., Li, C.W., Xia, W., Cha, J.H., Chan, L.C., Wu, Y., Chang, S.S., Lin, W.C., Hsu, J.M., Hsu, Y.H. et al. (2016). Deubiquitination and Stabilization of PD L1 by CSN5. Cancer Cell, 30, 925–939.

62. Lingaraju, G.M., Bunker, R.D., Cavadini, S., Hess, D., Hassiepen, U., Renatus, M., Fischer, E.S., and Thomä, N.H. (2014). Crystal structure of the human COP9 signalosome. Nature, 512, 161–165.

63. Liu, Y., Shah, S.V., Xiang, X., Wang, J., Deng, Z.B., Liu, C., Zhang, L., Wu, J., Edmonds, T., Jambor, C. et al. (2009). COP9-associated CSN5 regulates exosomal protein deubiquitination and sorting. Am J Pathol, 174, 1415–1425.

64. Liu, C., Yao, Z., Wang, J., Zhang, W., Yang, Y., Zhang, Y., Qu, X., Zhu, Y., Zou, J., Peng, S. et al. (2020). Macrophage-derived CCL5 facilitates immune escape of colorectal cancer cells via the p65/STAT3-CSN5-PD-L1 pathway. Cell Death Differ, 27, 1765–1781.

65. Llamas, E., Alirzayeva, H., Loureiro, R., and Vilchez, D. (2020). The intrinsic proteostasis network of stem cells. Curr Opin Cell Biol, 67, 46–55.

66. Lyapina, S., Cope, G., Shevchenko, A., Serino, G., Tsuge, T., Zhou, C., Wolf, D.A., Wei, N., Shevchenko, A., and Deshaies, R.J. (2001). Promotion of NEDD CUL1 conjugate cleavage by COP9 signalosome. Science, 292, 1382–1385.

67. Lykke-Andersen, K., Schaefer, L., Menon, S., Deng, X.W., Miller, J.B., and Wei, N. (2003). Disruption of COP9 signalosome Csn2 subunit in mice causes deficient cell proliferation, accumulation of p53 and cyclin E, and early embryonic death. Mol Cell Biol, 23, 6790–6797.

68. Ma, X.L., Xu, M., and Jiang, T. (2014). Crystal structure of the human CSN6 MPN domain. Biochem Biophys Res Commun, 453, 25–30.

69. Mantina, P., MacDonald, L., Kulaga, A., Zhao, L., and Hansen, D. (2009). A mutation in *teg-4*, which encodes a protein homologous to the SAP130 pre mRNA splicing factor, disrupts the balance between proliferation and differentiation in the *C. elegans* germ line. Mech Dev, 126, 417–429.

70. Marnik, E.A., Fuqua, J.H., Sharp, C.S., Rochester, J.D., Xu, E.L., Holbrook, S.E., and Updike, D.L. (2019). Germline maintenance through multifaceted activities of GLH/Vasa in *Caenorhabditis elegans* P granules. Genetics, 213, 923–939.

71. Merritt, C., and Seydoux, G. (2010). The Puf RNA-binding proteins FBF-1 and FBF-2 inhibit the expression of synaptonemal complex proteins in germline stem cells. Development, 137, 1787–1798.

72. Miller, R.K., Qadota, H., Stark, T.J., Mercer, K.B., Wortham, T.S., Anyanful, A., and Benian, G.M. (2009). CSN-5, a component of the COP9 signalosome complex, regulates the levels of UNC-96 and UNC-98, two components of M lines in *Caenorhabditis elegans* muscle. Mol Biol Cell, 20, 3608–3616.

73. Morgan, D. E., Crittenden, S. L., and Kimble, J. (2010). The *C. elegans* adult male germline: stem cells and sexual dimorphism. Dev Biol, 346, 204–214.

74. Morrison, S.J., Shah, N.M., and Anderson, D.J. (1997). Regulatory mechanisms in stem cell biology. Cell, 88, 287–298.

75. Oron, E., Mannervik, M., Rencus, S., Harari-Steinberg, O., Neuman-Silberberg, S., Segal, D., and Chamovitz, D.A. (2002). COP9 signalosome subunits 4 and 5 regulate multiple pleiotropic pathways in *Drosophila melanogaster*. Development, 129, 4399–4409.

76. Orsborn, A.M., Li, W., McEwen, T.J., Mizuno, T., Kuzmin, E., Matsumoto, K., and Bennett, K.L. (2007). GLH-1, the *C. elegans* P granule protein, is controlled by the JNK KGB-1 and by the COP9 subunit CSN-5. Development, 134, 3383– 3392.

77. Pan, L., Wang, S., Lu, T., Weng, C., Song, X., Park, J.K., Sun, J., Yang, Z., Yu, J., Tang, H., et al. (2014). Protein competition switches the function of COP9 from self-renewal to differentiation. Nature, 514, 233–236.

78. Pazdernik, N., and Schedl, T. (2013). Introduction to germ cell development in *Caenorhabditis elegans*. Adv Exp Med Biol, 757, 1–16.

79. Peth, A., Berndt, C., Henke, W., and Dubiel, W. (2007). Downregulation of COP9 signalosome subunits differentially affects the CSN complex and target protein stability. BMC Biochem, 8, 27.

80. Pfaffl, M.W. (2001). A new mathematical model for relative quantification in real time RT-PCR. Nucleic Acids Res, 29, e45.

81. Pintard, L., Kurz, T., Glaser, S., Willis, J.H., Peter, M., and Bowerman, B. (2003). Neddylation and deneddylation of CUL-3 is required to target MEI-1/Katanin for degradation at the meiosis-to-mitosis transition in *C. elegans*. Curr Biol, 13, 911– 921.

82. Prasad, A., Porter, D.F., Kroll-Conner, P.L., Mohanty, I., Ryan, A.R., Crittenden, S.L., Wickens, M., and Kimble, J. (2016). The PUF binding landscape in metazoan germ cells. RNA, 22, 1026–1043.

83. Puoti, A., and Kimble, J. (1999). The *Caenorhabditis elegans* sex determination gene *mog-1* encodes a member of the DEAH-Box protein family. Mol Cell Biol, 19, 2189–2197.

84. Puoti, A., and Kimble, J. (2000). The hermaphrodite sperm/oocyte switch requires the *Caenorhabditis elegans* homologs of PRP2 and PRP22. Proc Natl Acad Sci U S A, 97, 3276–3281.

85. Qin, N., Xu, D., Li, J., and Deng, X.W. (2020). COP9 signalosome: Discovery, conservation, activity, and function. J Integr Plant Biol, 62, 90–103.

86. Racher, H., and Hansen, D. (2012). PUF-8, a Pumilio homolog, inhibits the proliferative fate in the *Caenorhabditis elegans* germline. G3 (Bethesda), 2, 1197–1205.

87. Saha, A., and Deshaies, R.J. (2008). Multimodal activation of the ubiquitin ligase SCF by Nedd8 conjugation. Mol Cell, 32, 21–31.

88. Samuels, T.J., Järvelin, A.I., Ish-Horowicz, D., and Davis, I. (2020). Imp/IGF2BP levels modulate individual neural stem cell growth and division through. eLife, *9*, e51529.

89. Schindelin, J., Arganda-Carreras, I., Frise, E., Kaynig, V., Longair, M., Pietzsch, T., Preibisch, S., Rueden, C., Saalfeld, S., Schmid, B. et al. (2012). Fiji: an open source platform for biological-image analysis. Nat Methods, 9, 676–682.

90. Shackleford, T.J., and Claret, F.X. (2010). JAB1/CSN5: a new player in cell cycle control and cancer. Cell Div, 5, 26.

91. Sharon, M., Mao, H., Boeri Erba, E., Stephens, E., Zheng, N., and Robinson, C.V. (2009). Symmetrical modularity of the COP9 signalosome complex suggests its multifunctionality. Structure, 17, 31–40.

92. Shi, P., Zhang, J., Li, X., Li, W., Li, H., and Fu, P. (2021). Long non-coding RNA NORAD inhibition upregulates microRNA-323a-3p to suppress tumorigenesis and development of breast cancer through the PUM1/eIF2 axis. Cell Cycle, 20, 1295–1307.

93. Shin, H., Haupt, K.A., Kershner, A.M., Kroll-Conner, P., Wickens, M., and Kimble, J. (2017). SYGL-1 and LST-1 link niche signaling to PUF RNA repression for stem cell maintenance in *Caenorhabditis elegans*. PLoS Genet, 13, e1007121.

94. Sijen, T., Fleenor, J., Simmer, F., Thijssen, K.L., Parrish, S., Timmons, L., Plasterk, R.H., and Fire, A. (2001). On the role of RNA amplification in dsRNA triggered gene silencing. Cell, 107, 465–476.

95. Smialek, M.J., Ilaslan, E., Sajek, M.P., and Jaruzelska, J. (2021). Role of PUM RNA-binding proteins in cancer. Cancers (Basel), 13, 129.

96. Smith, P., Leung-Chiu, W.M., Montgomery, R., Orsborn, A., Kuznicki, K., Gressman-Coberly, E., Mutapcic, L., and Bennett, K. (2002). The GLH proteins, *Caenorhabditis elegans* P granule components, associate with CSN-5 and KGB-1, proteins necessary for fertility, and with ZYX-1, a predicted cytoskeletal protein. Dev Biol, 251, 333–347.

97. Söderberg, O., Leuchowius, K.J., Gullberg, M., Jarvius, M., Weibrecht, I., Larsson, L.G., and Landeregren, U. (2008). Characterizing proteins and their interactions in cells and tissues using the *in situ* proximity ligation assay. Methods, 45, 227–232.

98. Spassov, D.S., and Jurecic, R. (2002). Cloning and comparative sequence analysis of PUM1 and PUM2 genes, human members of the Pumilio family of RNA-binding proteins. Gene, 299, 195–204.

99. Spassov, D.S., and Jurecic, R. (2003). The PUF family of RNA-binding proteins, does evolutionarily conserved structure equal conserved function? IUBMB Life, 55, 359–366.

100. Spike, C.A., Tsukamoto, T., and Greenstein, D. (2022). Ubiquitin ligases and a processive proteasome protein clearance during the oocyte-to-embryo transition in *Caenorhabditis elegans*. Genetics, 221, iyac051.

101. Starostina, N.G., Lim, J., Schvarzstein, M., Wells, L., Spence, A.M., and Kipreos, E.T. (2007). A CUL-2 ubiquitin ligase containing three FEM proteins degrades TRA-1 to regulate *C. elegans* sex determination. Dev Cell, 13, 127–139.

102. Suh, N., Crittenden, S.L., Goldstrohm, A., Hook, B., Thompson, B., Wickens, M., and Kimble, J. (2009). FBF and its dual control of *gld-1* expression in the *Caenorhabditis elegans* germline. Genetics, 181, 1249–1260.

103. Tomoda, K., Kubota, Y., and Kato, J. (1999). Degradation of the cyclin dependent-kinase inhibitor p27Kip1 is instigated by Jab1. Nature, 398, 160–165.

104. Vilchez, D., Simic, M.S., and Dillin, A. (2014). Proteostasis and aging of stem cells. Trends Cell Biol, 24, 161–170.

105. Vogiatzoglou, A.P., Moretto, F., Makkou, M., Papamatheakis, J., and Kretsovali, A. (2022). Promyelocytic leukemia protein (PML) and stem cells: from cancer to pluripotency. Int J Dev Biol, 66, 85–95.

106. Voronina, E., and Greenstein, D. (2016). Germ cell fate determination in C. elegans. In eLS, John Wiley & Sons, Ltd: Chichester 1–8.

107. Voronina, E., and Seydoux, G. (2010). The *C. elegans* homolog of nucleoporin Nup98 is required for the integrity and function of germline P granules. Development, 137, 1441–1450.

108. Voronina, E., Paix, A., and Seydoux, G. (2012). The P granule component PGL-1 promotes the localization and silencing activity of the PUF protein FBF-2 in germline stem cells. Development, 139, 3732–3740.

109. Wang, C., Wilson-Berry, L., Schedl, T., and Hansen, D. (2012). TEG-1 CD2BP2 regulates stem cell proliferation and sex determination in the *C. elegans* germ line and physically interacts with the UAF-1 U2AF65 splicing factor. Dev Dyn, 241, 505–521.

110. Wang, X., and Voronina, E. (2020). Diverse roles of PUF proteins in germline stem and progenitor cell development in *C. elegans*. Front Cell Dev Biol, 8, 29.

111. Wang, X., Ellenbecker, M., Hickey, B., Day, N.J., Osterli, E., Terzo, M., and Voronina, E. (2020). Antagonistic control of *Caenorhabditis elegans* germline stem cell proliferation and differentiation by PUF proteins FBF-1and FBF-2. eLife, 9, e52788.

112. Wang, X., Olson, J.R., Rasoloson, D., Ellenbecker, M., Bailey, J., and Voronina, E. (2016). Dynein light chain DLC-1 promotes localization and function of the PUF protein FBF-2 in germline progenitor cells. Development, 143, 4643– 4653.

113. Wee, S., Geyer, R.K., Toda, T., and Wolf, D.A. (2005). CSN facilitates Cullin-RING ubiquitin ligase function by counteracting autocatalytic adapter instability. Nat Cell Biol, 7, 387–391.

114. Wei, N., and Deng, X.W. (1992). COP9: a new genetic locus involved in light-regulated development and gene expression in *Arabidopsis*. Plant Cell, 4, 1507–1518.

115. Wei, N., Chamovitz, D.A., and Deng, X.W. (1994). *Arabidopsis* COP9 is a component of a novel signaling complex mediating light control of development. Cell, 78, 117–124.

116. Wei, N., Serino, G., and Deng, X.W. (2008). The COP9 signalosome: more than a protease. Trends Biochem Sci, 33, 592–600.

117. Wickens, M., Bernstein, D.S., Kimble, J., and Parker, R. (2002). A PUF family portrait: 3’UTR regulation as a way of life. Trends Genet, 18, 150–157.

118. Wu, J.T., Lin, H.C., Hu, Y.C., and Chien, C.T. (2005). Neddylation and deneddylation regulate Cul1 and Cul3 protein accumulation. Nat Cell Biol, 7, 1014–1020.

119. Wu, Y., Deng, J., Rychahou, P.G., Qiu, S., Evers, B.M., and Zhou, B.P. (2009). Stabilization of snail by NF-kappaB is required for inflammation-induced cell migration and invasion. Cancer Cell, 15, 416–428.

120. Yang, X., Menon, S., Lykke-Andersen, K., Tsuge, T., Xiao, D., Wang, X., Rodriguez-Suarez, R.J., Zhang, H., and Wei, N. (2002). The COP9 signalosome inhibits p27(kip1) degradation and impedes G1-S phase progression via deneddylation of SCF Cul1. Curr Biol, 12, 667–672.

121. Yoon, S., Kawasaki, I., and Shim, Y.H. (2012). CDC-25.1 controls the rate of germline mitotic cell cycle by counteracting WEE-1.3 and by positively regulating CDK-1 in *Caenorhabditis elegans*. Cell Cycle, 11, 1354–1363.

122. Yoshida, A., Yoneda-Kato, N., and Kato, J.Y. (2013). CSN5 specifically interacts with CDK2 and controls senescence in a cytoplasmic cyclin E-mediated manner. Sci Rep, 3, 1054.

123. Yun, J., Tomida, A., Andoh, T., and Tsuruo, T. (2004). Interaction between glucose-regulated destruction domain if DNA topoisomerase IIα and MPN domain of Jab1/CSN5. J Biol Chem, 279, 31296–31303.

124. Zanetti, S., Meola, M., Bochud, A., and Puoti, A. (2011). Role of the *C. elegans* U2 snRNP protein MOG-2 in sex determination, meiosis, and splice site selection. Dev Biol, 354, 232–241.

125. Zhang, B., Gallegos, M., Puoti, A., Durkin, E., Fields, S., Kimble, J., and Wickens, M.P. (1997). A conserved RNA-binding protein that regulates sexual fates in the *C. elegans* hermaphrodite germ line. Nature, 390, 477–484.

126. Zhang, H., Gao, Z.Q., Wang, W.J., Liu, G.F., Shtykova, E.V., Xu, J.H., Li, L.F., Su, X.D., and Dong, Y.H. (2012). The crystal structure of the MPN domain from the COP9 signalosome subunit CSN6. FEBS Lett, 586, 1147–1153.

127. Zhou, C., Wee, S., Rhee, E., Naumann, M., Dubiel, W., and Wolf, D.A. (2003). Fission yeast COP9/signalosome suppresses cullin activity through recruitment of the deubiquitylating enzyme Ubp12p. Mol Cell, 11, 927–938.

