## Supplemental Figures and Tables for "COP9 signalosome component CSN-5 stabilizes PUF proteins FBF-1 and FBF-2 in *Caenorhabditis elegans* germline stem cells"

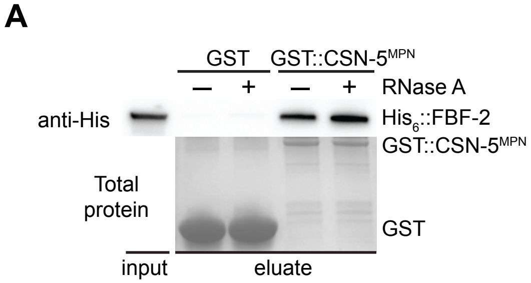


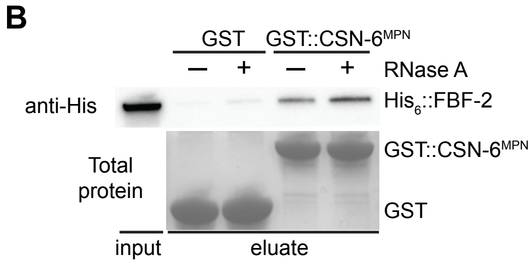


**Supplemental Figure 1.**

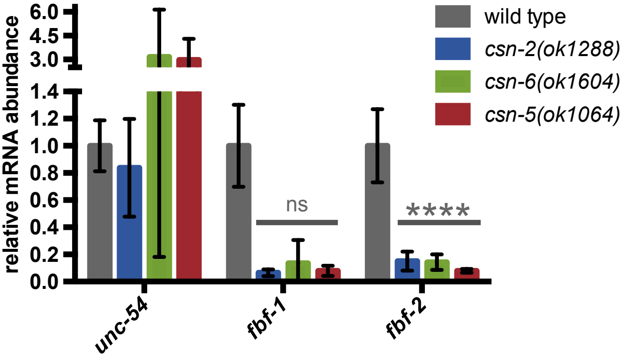


**Supplemental Figure 2.**

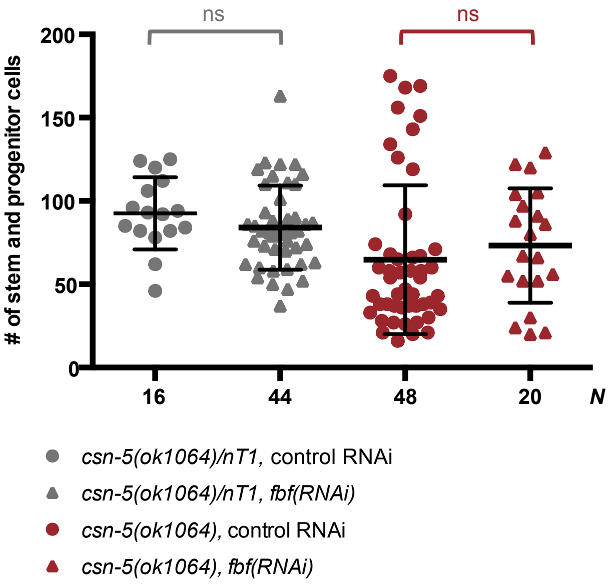


**Supplemental Figure 3.**

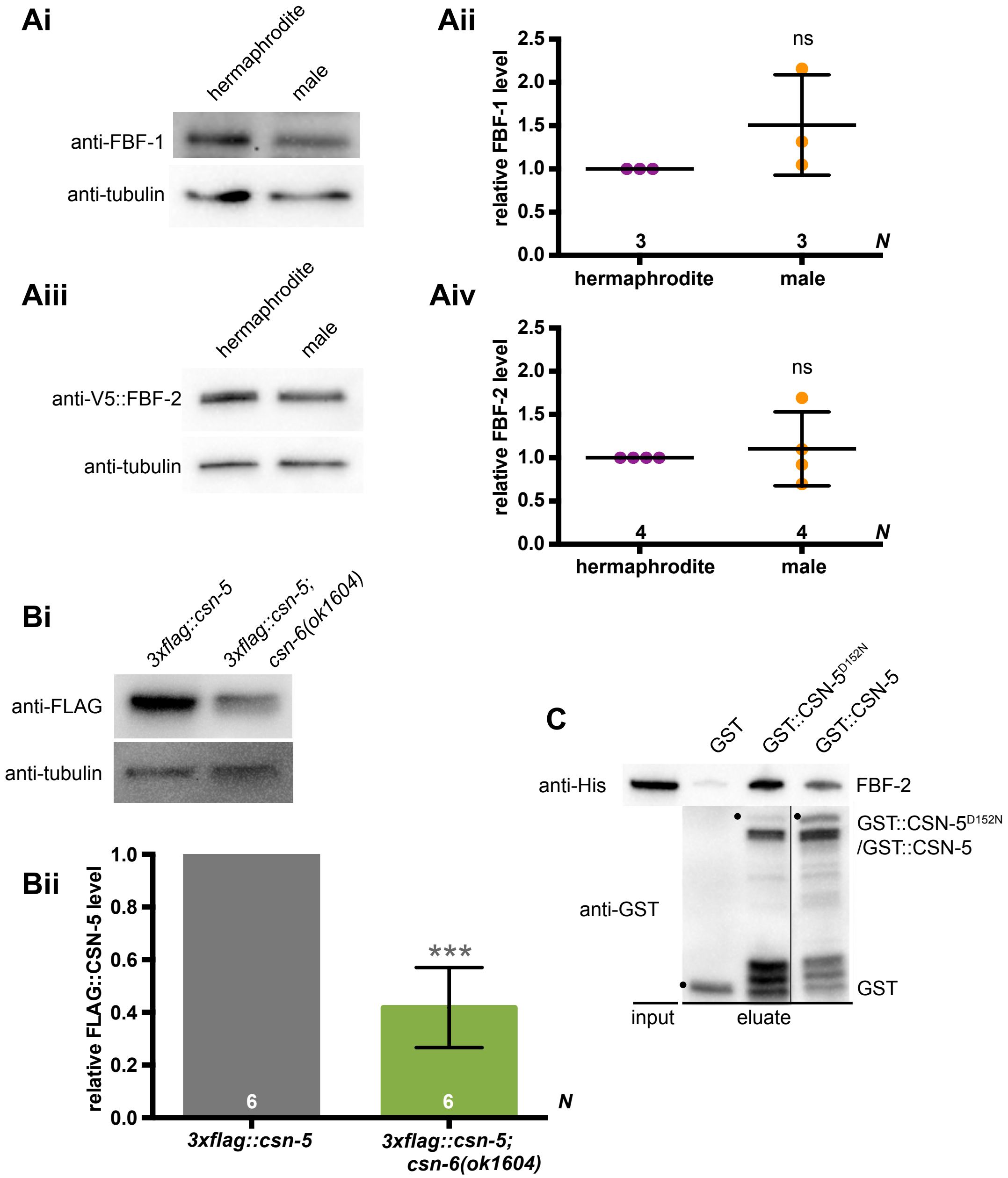


**Supplemental Figure 4.**

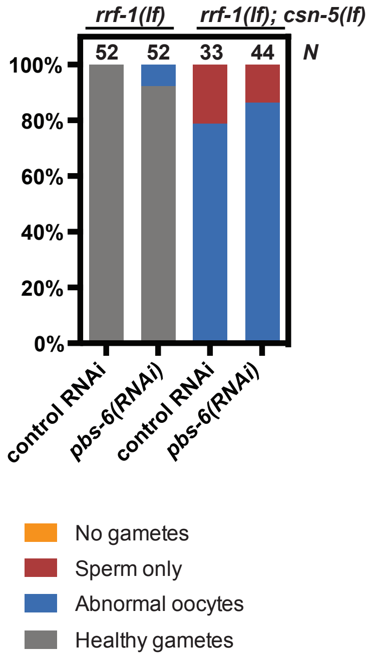


**Supplemental Figure 5.**

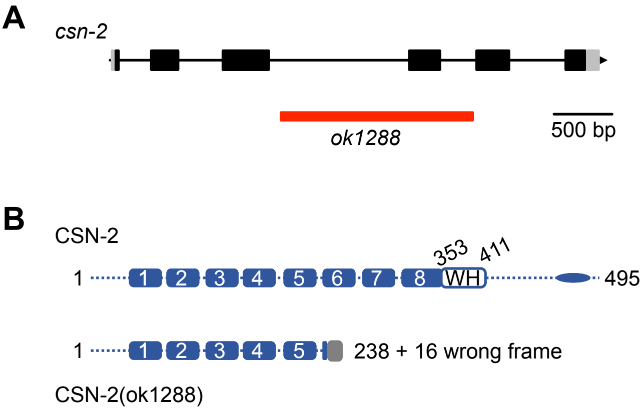


**Supplemental Figure 6.**

**Table S1. FBF-2 interactors identified via yeast two-hybrid assay**

Genes identified in a yeast two-hybrid screen with *fbf-2* as a bait, including a previously-documented interactor *sygl-1* (Shin et al., 2017).

| gene | function* | number of colonies | % positive |
|---|---|---|---|
| *ubh-4* | **UB**iquitin c-terminal **H**ydrolase, enables cysteine-type deubiquitinase activity | 130 | 48.87 |
| *csn-5* | **C**OP9 **S**ig**N**alosome subunit 5, catalytically active component of the complex responsible for deneddylation | 99 | 37.22 |
| *mlc-1* | **M**yosin **L**ight **C**hain component, part of muscle myosin complex | 12 | 4.51 |
| *vit-6* | **VIT**ellogenin structural genes (encoding egg yolk protein) | 4 | 1.50 |
| *tag-153* | predicted to be part of CCR4-NOT complex | 3 | 1.13 |
| *F48E3.4* | Predicted to be involved in proteolysis | 3 | 1.13 |
| *pas-4* | **P**roteasome **A**lpha **S**ubunit, predicted to be involved in protein degradation | 2 | 0.75 |
| *sygl-1* | **SY**nthetic **G**erm**L**ine proliferation defective, involved in regulation of germline stem cells, known interacting partner of FBF-2 (Shin et al., 2017) | 2 | 0.75 |
| *ddx-17* | **D**EA**D** bo**X** helicase homolog, predicted to enable RNA helicase activity | 1 | 0.38 |
| *mnat-1* | **MNAT** (menage a trois) TFIIH subunit, predicted to enable cyclin-dependent kinase activator activity | 1 | 0.38 |
| *lec-6* | Ga**LEC**tin, predicted to enable carbohydrate binding activity | 1 | 0.38 |
| *pas-2* | **P**roteasome **A**lpha **S**ubunit, predicted to be involved in protein degradation | 1 | 0.38 |
| *tdp-1* | **T**AR **D**NA-binding **P**rotein homolog, enables chromatin binding activity and single-stranded RNA binding activity | 1 | 0.38 |
| *egl-21* | **EG**g **L**aying defective, enables carboxypeptidase activity | 1 | 0.38 |
| *pud-2.2* | **P**rotein **U**p-regulated in **D**af-**2**(gf) | 1 | 0.38 |
| *fkb-2* | **FK**506-**B**inding protein family, predicted to enable isomerase activity and be involved in chaperone-mediated protein folding | 1 | 0.38 |

| | | | |
|---|---|---|---|
| *plc-4* | **P**hospho**L**ipase **C**, predicted to enable phosphatidylinositol phospholipase C activity | 1 | 0.38 |
| *K04C2.2* | Predicted to enable sequence-specific DNA binding activity and transcription corepressor activity | 1 | 0.38 |
| *R186.3* | Predicted to enable GTP binding activity | 1 | 0.38 |

*brief description of gene functions obtained from www.wormbase.org

### Table S2. Nematode strains used in this study.

| Genotype | Transgene description | Strain Number | Reference |
|---|---|---|---|
| wild-type | | N2 | Brenner, 1974 |
| *fbf-1(ok91)* | | JK3022 | Crittenden et al., 2002 |
| *fbf-2(q932)* | | JK5842 | Shin et al., 2017 |
| *csn-2(ok1288)/hT2* [*bli-4(e937) let-?(q782) qIs48*]*; fbf-2(q932)* | | UMT 438 | this study |
| *fbf-2(q932); csn-6(ok1604)/nT1* [*unc-?(n754) let-? qIs50*] | | UMT 465 | this study |
| *fbf-2(q932); csn-5(ok1064)/nT1*[*unc-?(n754) let-? qIs50*] | | UMT 452 | this study |
| *puf-3(q1058)* | | JK6080 | Haupt et al., 2020 |
| *csn-5(ok1064) puf-3(q1058)/nT1* [*unc-?(n754) let-? qIs50*] | | UMT 532 | this study |
| *csn-2(ok1288)/hT2* [*bli-4(e937) let-?(q782) qIs48*]*; fbf-1(ok91)* | | UMT 513 | this study |
| *rrf-1(pk1417); fbf-2(q932)* | | UMT 467 | this study |
| *rrf-1(pk1417); fbf-2(q932); csn-5(ok1064)/nT1*[*unc-?(n754) let-? qIs50*] | | UMT 478 | this study |
| *rrf-1(pk1417); fbf-2(q932); him-8(tm611)* | | UMT 468 | this study |
| **Transgenes: ORF + 3'UTR** | | | |
| *mntSi30 (pMT3.4; 3xFLAG::CSN-5) II; mntSi21 (pXW6.22; patcGFP) unc-119 (ed3) III* | *csn-5* prom::3xFLAG::CSN-5::*csn-5 3'UTR; gld-1* prom::patcGFP::*fbf-1 3'UTR* + *unc-119(+)* | UMT 463 | this study |
| *mntSi30 (pMT3.4; 3xFLAG::CSN-5) II; mntSi28 (pXW6.27; patcGFP::FBF-1) unc-119 (ed3) III* | *csn-5* prom::3xFLAG::CSN-5::*csn-5 3'UTR; gld-1* prom::patcGFP::FBF-1::*fbf-1 3'UTR* + *unc-119(+)* | UMT 462 | this study |
| *mntSi30 (pMT3.4; 3xFLAG::CSN-5) II; mntSi27 (pXW6.26; patcGFP::FBF-2) unc-119 (ed3) III* | *csn-5* prom::3xFLAG::CSN-5::*csn-5 3'UTR; gld-1* prom::patcGFP::FBF-2::*fbf-2 3'UTR* + *unc-119(+)* | UMT 461 | this study |
| *mntSi30 (pMT3.4; 3xFLAG::CSN-5) II* | *csn-5* prom::3xFLAG::CSN-5::*csn-5 3'UTR* | UMT 401 | this study |
| *mntSi30 (pMT3.4; 3xFLAG::CSN-5) II; csn-6(ok1604)/nT1*[*unc-?(n754) let-? qIs50*] | *csn-5* prom::3xFLAG::CSN-5::*csn-5 3'UTR* | UMT 464 | this study |
| *mntSi36 (pEO7.2; gld-1p::3xFLAG::CSN-5) II; csn-5(ok1064)/nT1 [unc-?(n754) let-? qIs50*] | *gld-1* prom:: 3xFLAG::CSN-5::*csn-5 3'UTR* | UMT 510 | this study |
| *mntSi38 (pEO7.4; gld-1p::3xFLAG::CSN-5(D152N)) II; csn-5(ok1064)/nT1 [unc-?(n754) let-? qIs50*] | *gld-1* prom:: 3xFLAG::CSN-5(D152N)::*csn-5 3'UTR* | UMT 514 | this study |

### Table S3. Antibodies used in this study.

| Primary Antibody | Host | Manufacturer; Cat. No. or reference | Dilution | Secondary Antibody | Host | Manufacturer; Cat. No. | Dilution |
|---|---|---|---|---|---|---|---|
| **Antibodies used for Western blot** | | | | | | | |
| Anti-myc | Mouse | DSHB; 9E10 | 1:1000 | Anti-mouse (IgG, light chain) | Goat | Jackson ImmunoResearch; 115-035-174 | 1:2000 |
| Anti-HA | Mouse | Thermo-Fisher; 26183 | 1:4000 | | | | |
| Anti-6xHis | Mouse | Sigma-Aldrich; H1029 | 1:2000 | Anti-mouse (IgG2a) | Goat | Southern Biotech; 10080-05 | 1:1000 |
| Anti-GST | Rabbit | Sigma-Aldrich; G7781 | 1:6000 | Anti-rabbit (IgG, H+L) | Goat | Jackson ImmunoResearch; 111-035-003 | 1:5000 |
| Anti-FBF-1 | Rabbit | Voronina et al., 2012; PA2388 | 1:10 | | | | |
| Anti-FLAG | Mouse | Sigma-Aldrich; F18094 | 1:1000 | Anti-mouse (IgG, H+L) | Goat | Jackson ImmunoResearch; 115-035-003 | 1:5000 |
| Anti-α-tubulin | Mouse | Sigma-Aldrich; T6199 | 1:500 | | | | |
| Anti- α-tubulin | Mouse | DSHB; AA4.4 | 1:4000 | | | | |
| Anti-V5 (IgG2a) | Mouse | Invitrogen; R96025 | 1:500 | | | | |
| **Antibodies for PLA** | | | | | | | |
| Anti-GFP (IgG) | Rabbit | Thermo-Fisher; G10362 | 1:40,000 | Anti-rabbit PLUS | Donkey | Sigma-Aldrich; DUO92002 | 1:5 |
| Anti-FLAG (IgG1) | Mouse | Sigma-Aldrich; F1804 | 1:1000 | Anti-mouse MINUS | Donkey | Sigma-Aldrich; DUO92004 | 1:5 |
| **Antibodies for immunostaining** | | | | | | | |
| Anti-FBF-1 | Rabbit | Voronina et al., 2012; PA2388 | 1:10 | Anti-rabbit (IgG) Alexa Fluor 488 | Goat | Jackson ImmunoResearch; 111-545-144 | 1:200 |
| Anti-REC-8 | Rabbit | Novus Biologicals; 29470002 | 1:500 | | | | |
| | | | | Anti-rabbit (IgG) Alexa Fluor 594 | Goat | Jackson ImmunoResearch; 111-585-144 | 1:500 |
| Anti-V5 (IgG2a) | Mouse | Invitrogen; R96025 | 1:500 | Anti-mouse (IgG) Alexa Fluor 488 | Goat | Invitrogen; A11029 | 1:200 |
| | | | | | Goat | | 1:500 |

| | | | | | | | |
|---|---|---|---|---|---|---|---|
| Anti-phospho-histone H3 | Mouse | Cell Signaling Technology; 9706L | 1:400 | Anti-mouse (IgG) Alexa Fluor 594 | | Jackson ImmunoResearch; 115-585-146 | |
| Anti-MSP | Mouse | DSHB; 4A5 | 0.2 µg/ml | Anti-mouse (IgG1) Alexa Fluor 488 | Goat | Jackson ImmunoResearch; 115-545-205 | 1:700 |

**Table S4. qPCR primer sequences.**

| Name | Primers (5' to 3') | Reference |
|---|---|---|
| act-1.qF | TGCTGATCGTATGCAGAAGC | Chauve et al., 2020 |
| act-1.qR | ATCCAGACGGAGTACTTGCG | Chauve et al., 2020 |
| unc-54.qF | AGAGAGCAGGTTTTGGAGGAT | Wang et al., 2020 |
| unc-54.qR | TTGAGGGTGACCTCATTTCC | Wang et al., 2020 |
| fbf-1.qF | GCTCTACCGAGATTGATCGTAA | Voronina and Seydoux, 2010 |
| fbf-1.qR | TTTGTCAACGGCAAACTTCATC | Voronina and Seydoux, 2010 |
| fbf-2.qF | TCTCATTCCAGGATTGGCTGA | Chauve et al., 2020 |
| fbf-2.qR | GAGTTGAACGGAGATTGGCA | Chauve et al., 2020 |
